# *TUBA1A* mutations identified in lissencephaly patients dominantly disrupt neuronal migration and impair dynein activity

**DOI:** 10.1101/377440

**Authors:** Jayne Aiken, Jeffrey K. Moore, Emily A. Bates

**Affiliations:** Department of Cell and Developmental Biology; University of Colorado School of Medicine; Aurora, CO, 80045; USA; Department of Pediatrics, University of Colorado School of Medicine; Aurora, CO, 80045; USA

**Keywords:** Tubulinopathy, Lissencephaly, Microtubules, Dynein, Neurodevelopment

## Abstract

‘Tubulinopathies’ are severe human brain malformations associated with mutations in tubulin genes. Despite the identification of many tubulin mutations in patients, we do not understand how these mutations impact the microtubule cytoskeleton, how the changes to microtubule function lead to brain malformations, or how different tubulin isotypes regulate microtubules to support normal neurodevelopment. *TUBA1A* α-tubulin is the most commonly affected tubulin isotype in tubulinopathy patients. Heterozygous mutations in *TUBA1A* have been identified in patients with diverse cortical malformations including microlissencephaly, lissencephaly, pachygyria, and polymicrogyria. Here we focus on mutations affecting the conserved arginine at position 402 (R402), which account for 30% of all reported *TUBA1A* mutations in patients. We demonstrate that exogenous expression of *TUBA1A*-R402C and *TUBA1A*-R402H patient alleles is sufficient to dominantly disrupt cortical neuron migration in the developing mouse brain, recapitulating the human lissencephaly phenotype. Intriguingly, ectopic expression of *TUBA1A*-R402C/H alleles does not alter morphology, axonal trafficking, or microtubule polymerization rates in cultured neurons, but does lead to subtle changes in axonal microtubule orientation. Further, we find that budding yeast α-tubulin with analogous R402C and R402H mutations assembles into microtubules but disrupts the activity of the microtubule motor dynein. The level of dynein impairment scales with abundance of R402 mutant α-tubulin in the cell. Together, our results support a model in which tubulinopathy mutations at R402 poison the microtubule network in young neurons by creating defective binding sites for dynein at the microtubule surface.

## INTRODUCTION

Tubulin genes have recently been implicated in severe and diverse neurodevelopmental defects in patients that can lead to intellectual deficits, epilepsy, and even fatality. These disorders, collectively termed ‘tubulinopathies’, result from dominant, *de novo*, missense mutations in tubulin genes, but how these alterations to tubulin proteins cause wide-ranging neuronal defects is largely unknown. The most commonly affected tubulin gene in tubulinopathy patients is *TUBA1A*, the human isoform of α1 tubulin that is selectively expressed in post-mitotic neurons, but not in neuronal progenitor or glia cells (Bahi-Buisson et al. 2014; Fallet-Bianco et al. 2014; Gloster et al. 1994; Coksaygan et al. 2006; Lewis et al. 1985; Gloster et al. 1999). α-tubulin protein forms a functional heterodimer with β-tubulin, providing the basic subunit that forms microtubules. Interestingly, patients with different *TUBA1A* mutations exhibit a wide spectrum of brain malformations (Keays et al. 2007; Poirier et al. 2007; Bahi-Buisson et al. 2008; Fallet-Bianco et al. 2008; Morris-Rosendahl et al. 2008; Kumar et al. 2010; Lecourtois et al. 2010; Poirier et al. 2013; Cushion et al. 2013; Bahi-Buisson et al. 2014). The variability of mutations and phenotypes underscores the possibility that different mutations may cause distinct defects in microtubule function, ultimately impacting brain development in drastically different ways (Bahi-Buisson et al. 2014; Aiken et al. 2017). However, we have little evidence of the molecular consequences of tubulin mutations, or how molecular changes to the microtubule network may give rise to the observed brain phenotypes.

Microtubules are long hollow polymers that dynamically grow and shrink to provide structure and generate force in a broad range of cellular contexts. In developing neurons, microtubules perform a variety of essential functions including neurite formation and outgrowth, neuronal migration, cargo transport, and synapse generation (Dent & Kalil 2001; Hu et al. 2012; Lin et al. 2012). Patient derived mutations can provide insight into how molecular level interactions impact each of these functions to ultimately dictate the stereotypical architecture of the brain. Many of the known *TUBA1A* mutations are associated with lissencephaly, such that patient brains exhibit fewer to absent cortical folds and disrupted cortical layering (Stratton et al. 1984; Poirier et al. 2007; Fallet-Bianco et al. 2008; Morris-Rosendahl et al. 2008; Kumar et al. 2010; Tian et al. 2010; Keays et al. 2007; Bahi-Buisson et al. 2014; Fallet-Bianco et al. 2014; Sohal et al. 2012). Lissencephaly is considered a neuronal migration disorder and is also associated with mutations that disrupt *LIS1*, which encodes a regulator of the microtubule-motor dynein, and *DCX*, which encodes the microtubule-associated protein (MAP) doublecortin (Fry et al. 2014). The association between these mutations and defects in neuronal migration suggest that appropriate neuronal migration relies on the microtubule/dynein interaction to generate the force required to move neurons to their ultimate positions (Tsai et al. 2007). Mechanistic studies corroborate the importance of dynein, microtubules, and their associated proteins LIS1 and doublecortin in neuronal migration and in the lissencephaly phenotype (Tanaka et al. 2004; Moon & Wynshaw-Boris 2014). Accordingly, migrating neurons may be uniquely sensitive to perturbations of the microtubule cytoskeleton.

Substitutions that affect R402 represent 30% of *TUBA1A* mutations identified in tubulinopathy, making this codon the most frequently mutated and suggesting this site is important for *TUBA1A’s* role in brain development. These mutations result in the substitution of the wild-type R402 residue to a cysteine (R402C), histidine (R402H), or leucine (R402L). R402C and R402H mutations are the most common, with only two reported cases of R402L (Morris-Rosendahl et al. 2008; Sohal et al. 2012). All three substitutions at R402 observed in patients cause moderate to severe lissencephaly phenotypes accompanied by corpus callosum defects, intellectual deficits, and occasionally microcephaly phenotypes (Poirier et al. 2007; Keays et al. 2007; Morris-Rosendahl et al. 2008; Kumar et al. 2010; Bahi-Buisson et al. 2014).

While many studies have linked *TUBA1A* mutations to severe human neurodevelopmental defects, little is understood about the mechanisms behind the diverse brain malformations that arise when *TUBA1A* mutations are present. Numerous, nearly identical α-tubulin isotypes exist in mammalian systems posing a challenge for investigating the role of *TUBA1A* in neurons. While post-mitotic neurons strongly and selectively express *TUBA1A* during development, a nearly identical isotype, *TUBA1B*, is expressed ubiquitously (Lewis et al. 1985; Miller et al. 1987). Genomic manipulation of a single α-tubulin isotype has presented an insurmountable challenge to date, as homology-directed repair techniques will target many α-tubulin genes, potentially disrupting α-tubulins that are required for cell division throughout development. Recently, tubulinopathy studies have employed alternative model systems to circumvent this challenge and investigate the molecular consequences of mutations to tubulin genes. For example, Tischfield *et al*. used budding yeast to interrogate the molecular mechanism behind *TUBB3* mutations that cause the ocular motility disorder CFEOM3 (Tischfield et al. 2010). Budding yeast provides a simplified eukaryotic system with only one β-tubulin and two α-tubulin genes. Tub1 is the major α-tubulin isotype expressed in yeast, comprising 90% of total α-tubulin (Schatz & Botstein 1986). In addition, yeast maintain many of the homologous microtubule associated proteins present in mammalian cells. Using yeast to investigate the functional consequences of mutations identified in tubulinopathy patients provides an opportunity to determine how a substitution alters tubulin function: 1) does the mutant α-tubulin form functional tubulin heterodimers, 2) can mutant heterodimers polymerize into microtubules, and/or 3) do mutant heterodimers alter binding of conserved microtubule associate proteins (MAPs) to the microtubule network (Aiken et al. 2017).

Here, we demonstrate that ectopic expression of *TUBA1A*-R402C and -R402H alleles are sufficient to disrupt cortical neuron migration in mouse, suggesting that these changes are responsible for the lissencephaly phenotype observed in patients. We then investigate the molecular mechanism responsible for the neuronal malformations caused by R402C and R402H mutants. While R402C/H mutants can dominantly disrupt cortical migration *in vivo*, they do not cause significant changes to neuronal morphology or microtubule polymerization in primary cortical neuron culture. Consistent with the neuronal migration phenotype, we reveal that R402C/H substitutions specifically disrupt the microtubule:dynein interaction and that the dynein phenotype scales with the amount of mutant tubulin expressed within the cell. Taken together, these findings suggest that R402 mutants cause lissencephaly by poisoning dynein activity in a unique setting when its function is crucial for neurodevelopment.

## RESULTS

### Mutations to conserved residue R402 dominantly disrupt cortical migration

Numerous studies have described the lissencephaly spectrum phenotype observed in tubulinopathy patients harboring heterozygous *TUBA1A*-R402C and R402H mutations (Keays et al. 2007; Poirier et al. 2007; Morris-Rosendahl et al. 2008; Kumar et al. 2010; Fallet-Bianco et al. 2008; Bahi-Buisson et al. 2014; Fallet-Bianco et al. 2014; Kamiya et al. 2014), but whether these mutations alone are sufficient to dominantly disrupt cortical migration has not been established. To examine how *TUBA1A*-R402C and *TUBA1A-R402H* impact neuronal migration, we performed *in utero* electroporation to introduce ectopic *TUBA1A* expression in the ventricular zone of C57Bl6 wild-type mice (Figure 1). Two control plasmids express either cytoplasmic GFP alone (empty vector), or GFP along with wild-type *TUBA1A*. Two experimental constructs express GFP with either *TUBA1A-R402C* or *TUBA1A*-R402H. Over-expression of the wild-type *TUBA1A* caused no alteration to migration compared to the empty vector control. In contrast, overexpression of either *TUBA1A*-R402C or *TUBA1A*-R402H alleles lead to a marked perturbation of cortical neuron migration, with only 24.3 ± 1.7% (mean ± SEM) and 27.6 ± 2.5% percent of total GFP signal present in the cortical plate, respectively, compared to 56.7 ± 3.1% of the empty vector and 56.8 ± 3.2% for wild-type *Tuba1a* expression controls (Figure 1A-1C; p-value<0.0001, t-test). In all conditions, including *TUBA1A*-R402C/H mutant expression, we observed neurons with long axons (Figure S1A) and visible leading processes (Figures S1B and S1C) based on cytosolic GFP, suggesting that the migration failure is not accompanied by failure to send out axons or leading processes. Further, we show that while neurons expressing mutant *TUBA1A* do not appropriately migrate, non-electroporated neurons in the same sections reach upper cortical layers unperturbed, as shown by *Cux1* staining (Figure 1D). This suggests that the migration failure is intrinsic to the transfected neurons ectopically expressing the *TUBA1A*-R402C/H mutant alleles and not attributable to disruption of the radial glial cells.

**Figure 1.**
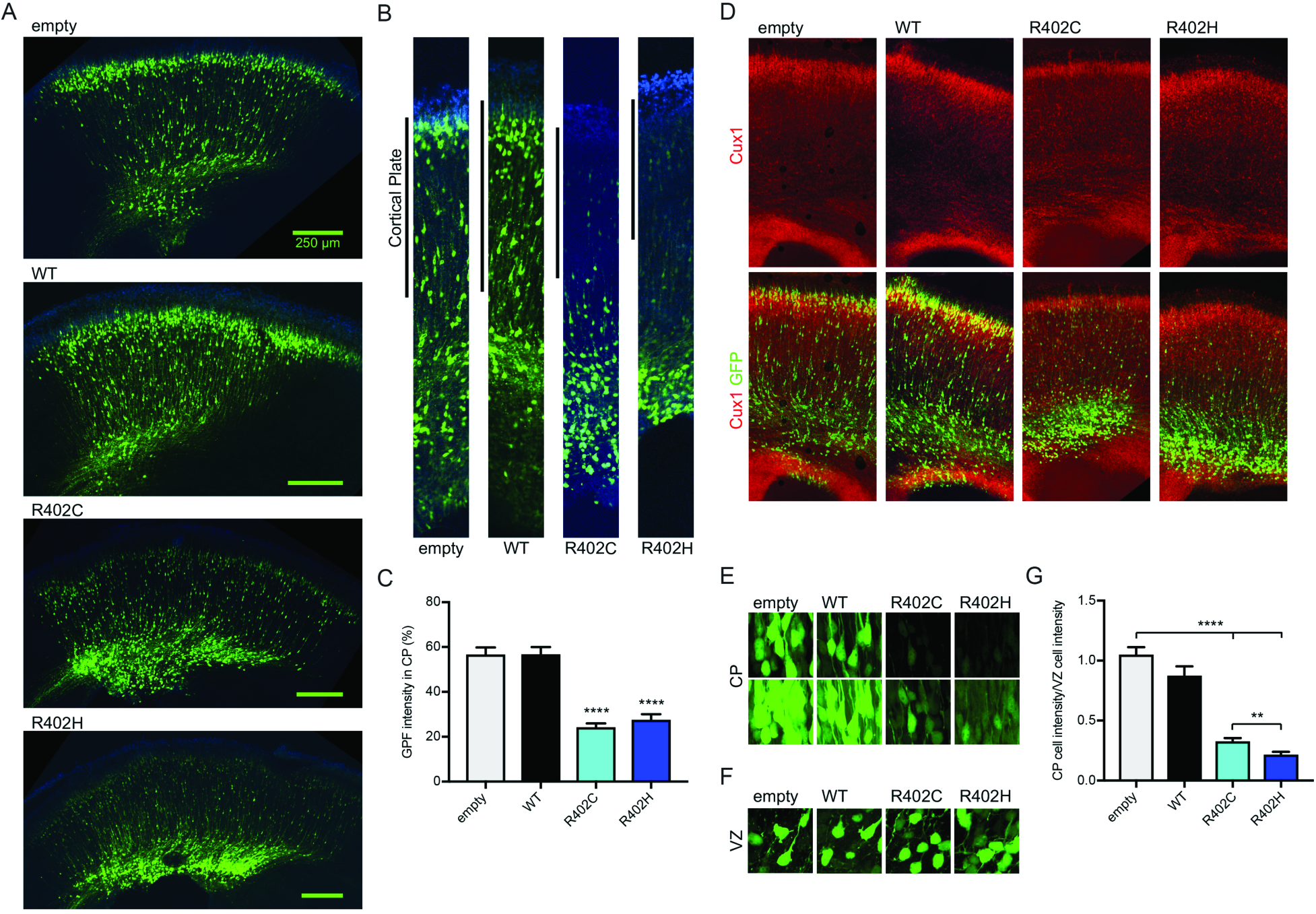
*TUBA1A*-R402C/H mutants dominantly disrupt neuronal migration in the developing mouse cortex. (A) Coronal sections from E18.5 mouse brain electroporated at E14.5 with pCIG2 vectors: empty vector (empty), wild-type *TUBA1A* (WT), *TUBA1A*-R402C (R402C), or *TUBA1A*-R402H (R402H). (B) Representative regions of cortex analyzed for cortical plate fluorescence. (C) Percentage of GFP signal in the cortical plate. For each condition, three coronal sections from at least four separate animals were analyzed. Data are represented as mean ± SEM. Quadruple asterisks indicate significant difference compared to wild type, by *t*-test (*p*<0.0001). (D) Cux1 staining of electroporated sections reveals position of upper layer cortical neurons. (E) Representative images of cortical plate neurons for each condition, at two different exposures. (F) Representative images ventricular zone neurons for each condition. (G) Quantification of mean GFP intensity of cortical plate neurons normalized GFP intensity of ventricular zone neurons. Data are represented as mean ± SEM. At least 45 neurons for each condition were measured (ROI=200 μm^2^). Quadruple asterisks indicate significant difference compared to wild.

Intriguingly, the presence of neurons containing low cytosolic GFP fluorescence intensity in the cortical plate of *TUBA1A*-R402C/H electroporated sections suggests that neurons with lower levels of mutant expression may migrate properly. For empty vector control, cells in the cortical plate are as intense as cells in the ventricular zone, with a normalized intensity ratio of 1.05 ± 0.06 (mean GFP intensity per cortical plate cell/mean GFP intensity per ventricular zone cell ± SEM). *TUBA1A* wild-type controls also contain similar GFP intensities in the cortical plate and ventricular zone, with a normalized intensity ratio of 0.88 ± 0.077 (p-value=0.0866, compared to empty vector; t-test). However, the *TUBA1A*-R402C/H-expressing neurons that reach the cortical plate exhibit significantly lower cytosolic GFP fluorescence with normalized intensity ratios of 0.33 ± 0.027 and 0.22 ± 0.024, respectively (p-value<0.0001, compared to empty vector; Figure 1E-1G and Figure S1B). This suggests that only neurons expressing lower levels of *TUBA1A*-R402C or *TUBA1A*-R402H escape the ventricular zone to populate the cortical plate. These data show that expression of *TUBA1A*-R402C and *TUBA1A*-R402H alleles dominantly disrupt cortical neuron migration, and the severity of the phenotype may depend on the level of *TUBA1A*-R402C/H expression in the cell.

### Ectopic expression of *TUBA1A* mutants is not sufficient to disrupt neuronal function in primary cortical culture

The developmental period when cortical neurons are born and migrate involves several microtubule-dependent functions: forming neurites, polarizing to contain one axon and the appropriate number of dendrites, trafficking cargoes to growth cones, and migrating along radial glial cells to reach the correct cortical layer. We set out to discover whether defects in these microtubule-dependent functions might underlie the cortical migration defects observed with the expression of *TUBA1A-R402C/H* alleles.

We first interrogated how ectopic expression of *TUBA1A* mutant alleles impact neuronal microtubule polymerization. For these experiments, we expressed *TUBA1A*-R402C or R402H constructs, *TUBA1A* wild-type constructs, or empty vectors in primary rat cortical neurons. In addition to our mutant alleles of interest, *TUBA1A*-R402C and -R402H, we also assessed the impact of *TUBA1A*-P263T mutant constructs, which had previously been suggested to decrease microtubule polymerization rate in COS7 and primary rat cortical neurites (Tian et al. 2010). To visualize microtubule polymerization events, we constructed pCIG2 vectors that co-express the *TUBA1A* alleles with GFP-MACF43. MACF43 binds to EB proteins and tracks the plus-ends of growing microtubules (Yau et al. 2016). Using this relatively inert plus-end marker removes the complication of overexpressing labeled EB proteins that act as microtubule polymerization factors and can impact polymerization rates (Vitre et al. 2008). We introduced these constructs via nucleofection in dissociated neurons before plating, and performed experiments to assess the effect of ectopic *TUBA1A* mutant alleles on axonal microtubule dynamics. For consistency in this analysis, we measured microtubule polymerization in proximal axons of DIV11 neurons (50-200 μm from the soma), as determined by staining with axon initial segment (AIS) marker external neurofascin (extNF) (Figure 2A). Using at least 17 cells from a minimum of three separate nucleofections, and analyzing 141-452 individual polymerization events for each condition, we determined that that ectopic expression of *TUBA1A*-R402C, -R402H or -P263T mutants does not significantly alter microtubule polymerization rates (Figure 2C and Table S1). Throughout this experiment, we noticed a large variability in microtubule polymerization rates from cell to cell, which often varied with different primary neuron nucleofections. For example, for each condition (empty vector, WT control, P263T, R402C, or R402H) we observed cells containing slowly polymerizing microtubules and other cells containing quickly polymerizing microtubules (Figure 2B).

**Figure 2.**
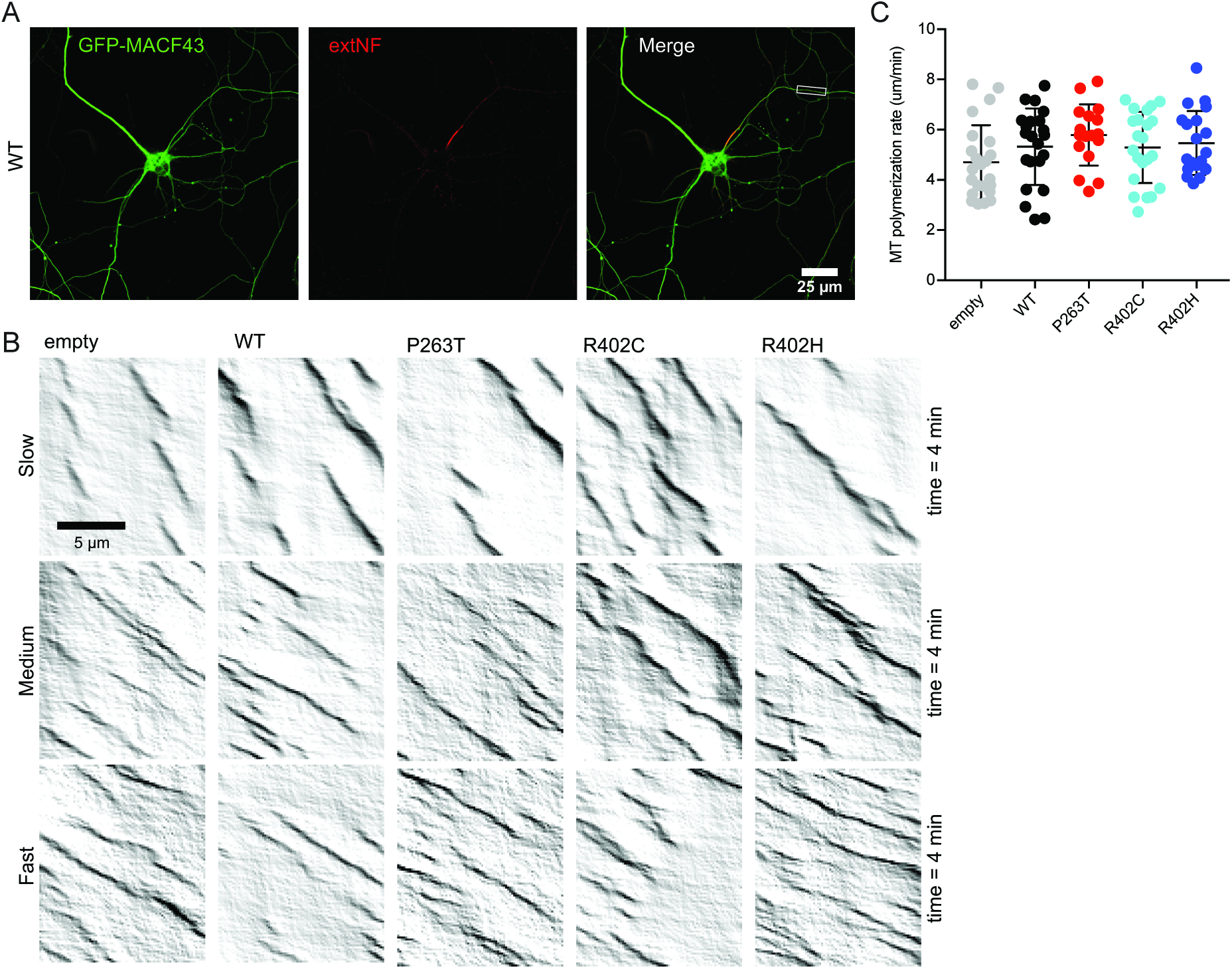
Ectopic expression of TUBA1A mutant alleles is not sufficient to disrupt axonal microtubule polymerization in primary neuronal culture. (A) Representative image of DIV11 primary rat cortical neuron expressing pCIG2-Tuba1a(WT)-ires-GFP-Macf43 and live-stained with extNF to reveal AIS. Inset reveals axonal segment selected for kymograph analysis. (B) Representative kymographs from each condition displaying variation in microtubule polymerization rates observed across cells. (C) Microtubule polymerization rate. Each data point represents cellular mean microtubule polymerization rate, with bars displaying mean ± SEM. No significant differences to empty vector or WT control, with significance determined as p<0.05.

In addition to microtubule polymerization, we also tested the effect of the *TUBA1A*-R402C/H mutant alleles on neurite outgrowth and polarization. Here again no significant difference was observed, with all conditions producing a similar number of neurites that appropriately extend and polarize (Figure S2 and Table S2). Finally, we measured axonal trafficking in neurons ectopically expressing *TUBA1A*-R402C/H alleles. For these experiments, we constructed pCIG2 plasmids that co-express *TUBA1A* alleles with the lysosomal marker Lyso-20-mNeonGreen and identified the axon using AIS marker extNF (Figure S3A). No significant difference was detected in any trafficking parameter measured (Figure S3 and Table S3). Given these results, we conclude that ectopic expression of *TUBA1A*-R402C, -R402H, or -P263T is not sufficient to disrupt the axonal microtubule network in this rat primary cortical neuron culture system.

### R402C/H mutations may disrupt the polarity of axonal microtubules

In addition to microtubule polymerization rate data, our axonal GFP-MACF43 experiments allowed us to determine the polarity of microtubules polymerizing in the axon. Normally, axonal microtubules are uniformly oriented with plus-ends toward the distal axon (Baas et al. 1988). We used our live imaging of neurons stained with extNF to ask whether axonal microtubule polarity is disrupted by *TUBA1A*-R402C/H mutants (Figure S4). MACF43 comets that project out from the soma to the distal axon represent anterograde oriented microtubules, while MACF43 comets in the opposite direction, from distal axon to soma, indicate the presence of microtubules oriented in the retrograde direction (Figure S4A). While microtubule density and polymerization rate are not significantly altered in neurons ectopically expressing *TUBA1A*-R402C/H mutants (Figure 2C and S4B), we did detect a small but significant change in the percentage of retrograde axonal microtubule polymerization events, comprising 6.92 ± 1.10% of all polymerizing axonal microtubules in R402C and 6.46 ± 1.07% in R402H, compared to 3.73 ± 0.86% in neurons containing empty vector (p=0.0251 for *TUBA1A*-R402C and p=0.0387 for *TUBA1A*-R402H, by Chi-square test; Figure S4C). Neurons expressing wild-type *TUBA1A* exhibited 5.09 ± 0.992% retrograde polymerization events, which is not significantly different than either empty vector controls or the *TUBA1A*-R402C/H mutants (Figure S4C). We also analyzed the locations of retrograde microtubule polymerization along the axon, and found that these exhibit no obvious sites of enrichment along the first ~150 microns of the axon (Figure S4D). These results suggest that while *TUBA1A*-R402C/H mutants do not drastically alter rates of microtubule polymerization, they may disrupt the polarized organization of microtubules in the axon.

### Mutations mimicking R402C/H in yeast α-tubulin lead to polymerization competent heterodimers

In principle, mutations to *TUBA1A* could alter α-tubulin function at the levels of: 1) tubulin heterodimer formation (Figure 3A), 2) tubulin heterodimer polymerization to form microtubules (Figure 3B), or 3) MAP binding to polymerized microtubules (Figure 3C), or a combination of these effects (Aiken et al. 2017). To isolate the functional consequences of *TUBA1A*-R402C and R402H substitutions on a molecular level, we made analogous mutations in budding yeast α-tubulin. Budding yeast is an ideal system to test specific molecular hypotheses about microtubule function because they contain fewer tubulin isotypes, mutations can be made at the endogenous chromosomal locus, and single microtubules can be visualized by light microscopy. We introduced the mutations to the major yeast α-tubulin isotype *TUB1* at the codon analogous to R402, generating *tub1*-R403C and *tub1*-R403H mutant stains. We first tested whether R403 mutants alter Tub1 protein levels by performing western blots on lysates from the integrated α-tubulin mutant yeast strains and probed with pan α-tubulin antibody, and found no significant change to total α-tubulin levels in either mutant strain compared to wild type (Figure 3D).

**Figure 3.**
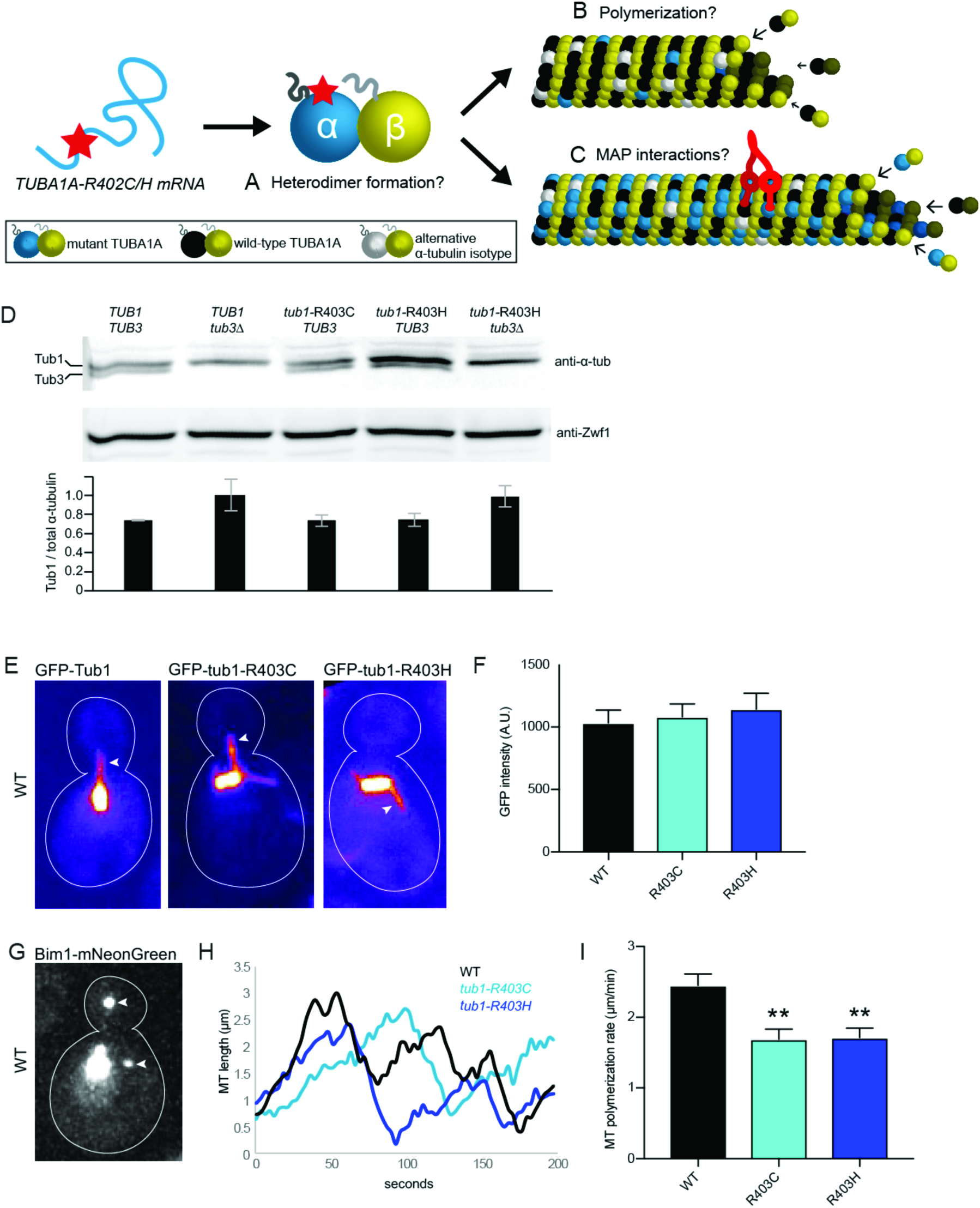
α-tubulin R402C/H mutants form polymerization competent tubulin heterodimers in *S. cerevisiae*. (A-C) Possible consequences of α-tubulin R402C/H mutants on tubulin function. R402C/H mutants may disrupt tubulin heterodimer formation (A), polymerization into microtubule lattice (B), and/or microtubule-associated-protein (MAP) interactions. (D) Western blot of α-tubulin protein in lysates from the indicated strains. Blots were also probed for Zwf1/G6PD as a loading control. Intensity of α-tubulin bands from 3 separate experiments, normalized to Zwf1 loading control. Data are represented as mean ± SeM. Strains: wild type, yJM596; tub3Δ, yJM0103; tub1-R403C, yJM2240, yJM2245; tub1-R403H, yJM2121, yJM2263; tub1-R403H tub3Δ, yJM2533. (E) Images of microtubules labeled with GFP-Tub1, GFP-tub1-R403C, or GFP-tub1-R403H. Arrows point to astral microtubules. (F) Quantification of GFP signal per micron of astral microtubule. 29 microtubules were measured for each strain. Data are represented as mean ± SEM. Strains: wild type GFP-TUB1, yJM1237, yJM1887, yJM0562; GFP-tub1-R403C, yJM1872, yJM2112, yJM2113; GFP-tub1-R403H, yJM1873, yJM2114, yJM2115. (G) Image of a wild-type cell expressing Bim1-mNeonGreen. Arrows point to astral microtubule plus ends. (H) Representative life plots of astral microtubule dynamics in wild type, tub1-R403C, and tub1-R403H mutants. Astral microtubule length was measured over time as the distance between Bim1-mNeonGreen at the microtubule plus end and the minus end at the proximal spindle pole. (I) Mean polymerization rates. Data are represented as mean ± SEM. Double asterisks indicate significant difference compared to wild type, by *t*-test (*p*<0.01). Strains: wild type, yJM2188, yJM2189; tub1-R403C, yJM2190, yJM2191; tub1-R403H, yJM2192, yJM2193.

In yeast, cell growth assays can be used as a measurement of the functionality of mutant tubulin. To further assess the competency of tub1-R403C/H, we tested these mutants for sensitivity to microtubule the destabilizing drug, benomyl. We performed growth assays using heterozygous diploid yeast, which contain one copy of *tub1*-R403C/H and one copy of wild-type *TUB1*. We also used haploid yeast strains that contain a single copy of either the mutant *tub1*-R403C/H or wild-type *TUB1* controls. The *tub1*-R403H mutant exhibits a level of growth and benomyl sensitivity that is similar to wild-type controls in both the diploid and haploid cells (Figure S5A and S5B). This indicates that the R403H mutant can grossly rescue α-tubulin function in yeast. In contrast, the *tub1*-R403C mutant exhibits hypersensitivity to benomyl in both the haploid and heterozygous diploid cells (Figure S5A and S5B). The growth disruption caused by the tub1-R403C mutant in heterozygous diploid cells is similar to *tub1*Δ/*TUB1* heterozygous null diploids, where one copy of *TUB1* is ablated. These data suggest that R403C and R403H substitutions may lead to differing effects on microtubule function.

To determine whether *tub1*-R403C/H is sufficient to support α-tubulin function, we tested for viability in the absence of the other yeast α-tubulin isotype, *TUB3*. Here, we again observed a difference between the *tub1*-R403C and *tub1*-R403H. While we were able to generate a *tub1*-R403H *tub3*Δ double mutant haploid strain, *tub1*-R403C is lethal when combined with the *tub3*Δ knockout (Table S5). This additional evidence suggests that *tub1*-R403C and *tub1*-R403H substitutions do not lead to the same functional consequences; the cysteine substitution may be more detrimental.

We next tested whether the R403 mutants alter tubulin assembly activity in two separate assays: first by lattice incorporation and second by measuring microtubule dynamics. We tested whether the mutant protein could assemble into microtubules by measuring the abundance of GFP-labeled wild-type Tub1, tub1-R403C, or tub1-R403H present in single astral microtubules in yeast cells. Importantly, these cells also express wild-type, unlabeled Tub1 and Tub3 α-tubulin; therefore, the level of GFP signal per unit length of microtubule reflects a competition between GFP-labeled and unlabeled α-tubulin for assembly into the lattice. This competition analysis revealed that mutant GFP-tub1-R403C and GFP-tub1-R403H incorporate into microtubules at levels similar to wild-type GFP-Tub1, as the fluorescence intensity of single microtubules measured in mutant strains was indistinguishable from wild-type strains (R403C p=0.73, R403H p=0.50 by t-test; Figure 3E and 3F). These data indicate that tub1-R403C and tub1-R403H support normal levels of heterodimer formation and microtubule assembly. To determine how mutant R403C/H α-tubulin affects microtubule dynamics, we crossed *tub1*-R403 mutants with a strain that also expresses fluorescently labeled Bim1, the yeast homologue of microtubule plus-end binding protein EB. Bim1 binds to the plus ends of microtubules and allows visualization of both polymerization and depolymerization events (Figure 3G and 3H). This analysis revealed that expression of both *tub1*-R403C and *tub1*-R403H lead to decreased microtubule polymerization rates of 1.69 ± 0.82 μm/minute (mean ± SEM) and 1.71 ± 0.73 μm/minute, respectively, compared to the wild-type polymerization rate of 2.45 ± 1.1 μm/minute (p=0.0014 for R403C, p=0.0018 for R403H, by t-test; Figure 3H and 3I, Table S4). Therefore, our results indicate that while *tub1*-R403C/H mutants are competent at assembling into microtubules, they polymerize more slowly than wild-type Tub1.

### *tub1*-R403C/H mutations perturb dynein function in yeast

A recent study used directed mutagenesis to demonstrate that changing arginine to alanine at position 403 of Tub1 disrupts dynein binding to microtubules (Uchimura et al. 2015). To investigate whether the histidine and cysteine substitutions similarly perturb dynein, we analyzed the activity of dynein in *tub1*-R403H or *tub1*-R403C mutant yeast. Unlike in neurons, where dynein performs many important tasks, yeast dynein performs the single, important role of generating pulling forces on astral microtubules to move the mitotic spindle across the nascent plane of cytokinesis.

To determine how *tub1*-R403C/H mutants affect dynein activity, we imaged mitotic spindle movement in strains simultaneously expressing GFP-*tub1*-R403C/H mutant and unlabeled *tub1*-R403C/H from the endogenous locus, so that all Tub1 protein in the cell contains the mutation. To isolate dynein-mediated spindle sliding events and prevent spindle elongation at anaphase, we treated the cells with hydroxyurea to arrest in S-phase. Under these conditions, dynein moves the spindle by promoting ‘sliding events’, where growing astral microtubules collide with the cell cortex, dynein attaches to a receptor at the cortex, and dynein motor activity slides the microtubule and attached spindle toward the anchored motor (Figure 4A; (Estrem et al. 2017; Carminati & Stearns 1997). The frequency of sliding events was calculated as the number of sliding events divided by the total number of microtubule-cortex collisions observed in the 10-minute imaging period. The frequency of dynein sliding events decreased from 13.9 ± 2.39% in wild-type cells to 6.47 ± 1.24% in *tub1*-R403C and 5.74 ± 1.07% in R403H (p=0.0042 for R403C, p=0.0012 for R403H, by t-test; Figure 3B and 3C). When sliding events did occur in the mutant cells, the distance moved by the mitotic spindle was significantly decreased, from the wild-type average sliding distance of 1.15 ± 0.08 μm to 0.73 ± 0.15 μm in *tub1*-R403C and 0.77 ± 0.09 μm in R403H, respectively (p=0.0078 for R403C, p=0.0047 for R403H, by t-test; Figure 3D). These data indicate that when *tub1*-R403C/H mutant microtubules hit the cortex, they rarely initiate a dynein sliding event. Additionally, when a rare sliding event does occur, the presence of *tub1*-R402C/H mutants limits the extent of spindle movement. Therefore, dynein function is diminished in *tub1*-R403C/H mutants, but not completely ablated.

**Figure 4.**
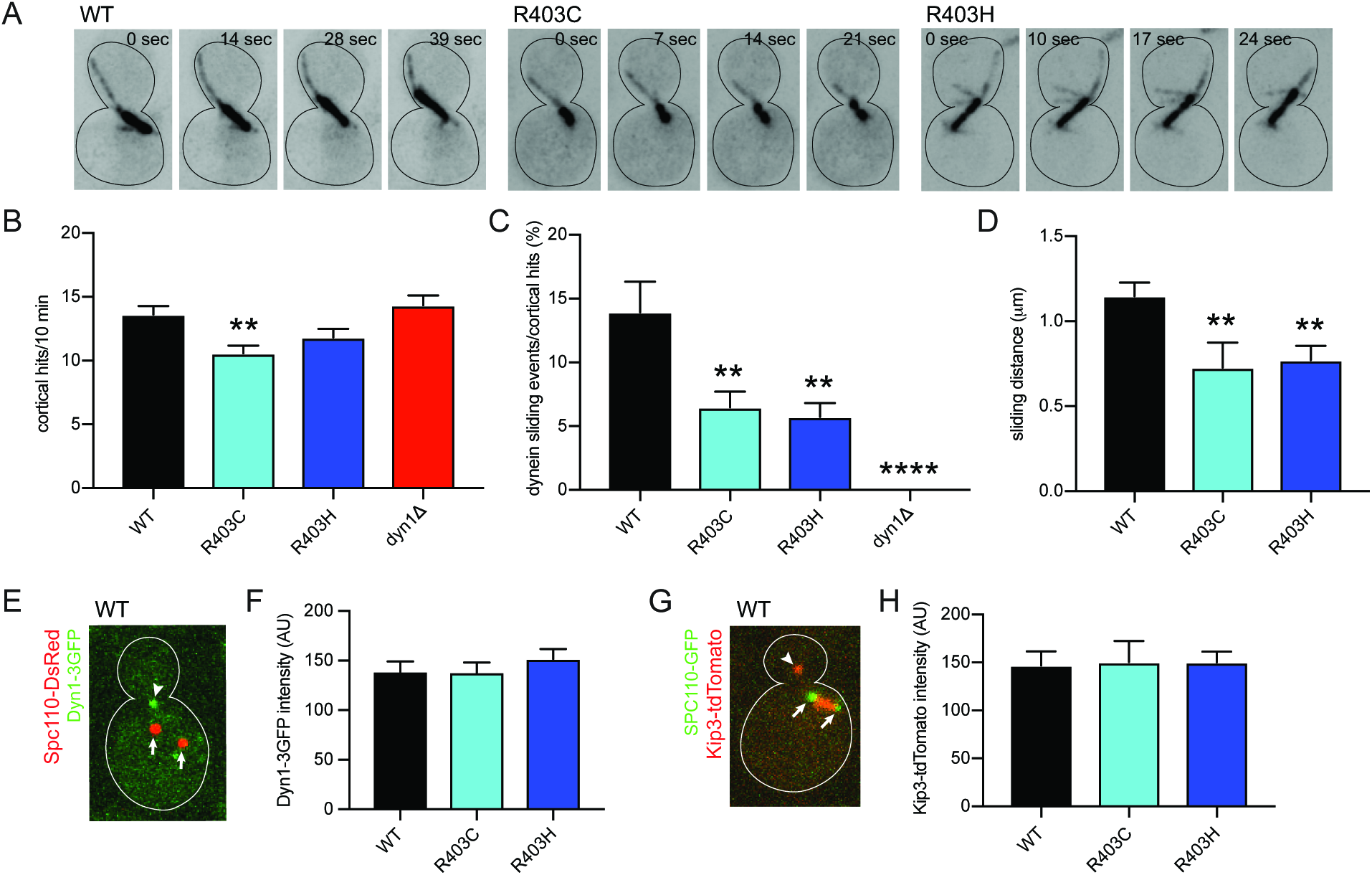
α-tubulin R402C/H mutants disrupt dynein activity in *S. cerevisiae*. (A) Time-lapse images of cells labeled with GFP-tub1, GFP-tub1-R403C, or GFP-tub1-R403H to illustrate representative dynein sliding events. Dynein sliding is defined by spindle translocation initiated by a microtubule-cortex interaction. (B) Quantification of microtubule-cortical hits in a 10 minute period. (C) Percent of microtubule-cortex hits that become dynein sliding events. (D) Quantification of average sliding distance, measured by spindle translocation during sliding event (μm). At least 34 cells were analyzed for each strain. Data are represented as mean ± SEM. Double asterisks indicate significant difference compared to wild type, by *t*-test *p*<0.01). Quadruple asterisks indicate significant difference compared to wild type, by *t*-test (*p*<0.0001). Strains: wild type, yJM1887; dyn1Δ, 1023; tub1-R402C, yJM2216; tub1-R402H, yJM2218. (E) Image of a wild-type cell expressing Dyn1-3GFP and spindle pole marker Spc110-DsRed. Arrowhead points to dynein localized to astral microtubule plus end. Arrows point to spindle pole bodies. (F) Quantification of GFP signal at microtubule plus ends. At least 39 dynein-plus end foci were measured for each strain. Error bars are SEM. Strains: wild type, yJM0307, yJM0308; *tub1*-R403C, yJM2202, yJM2203; tub1-R403H, yJM2204. (G) Image of a wild-type cell expressing Kip3-tdTomato and Spc110-GFP. Arrowhead points to Kip3 localized astral microtubule plus end. Arrows point to spindle pole bodies. (H) Quantification of tdTomato signal at plus ends. 20 Kip3-plus end foci were measured for each strain. Data are represented as mean ± SEM. Strains: wild type, yJM2696, yJM2716; tub1-R403C, yJM2698, yJM2718; tub1-R403H, yJM2697, yJM2717.

We also assessed dynein function using a genetic assay. When dynein function is disrupted, yeast cells position the mitotic spindle through a compensatory pathway involving Bim1 and actin-trafficking. Therefore, yeast cells can divide when one pathway is compromised, but are inhibited when both pathways are compromised. To genetically test whether the *tub1*-R403C/H mutants are acting to disrupt the dynein pathway, we generated double mutant strains combining *tub1*-R403C/H mutants with *dyn1*Δ or *bim1*Δ null mutations by meiotic cross (Table S5). Double mutants combining *tub1*-R403C or R403H mutations with *bim1*Δ exhibit significantly impaired growth (Figure S5E and S5F), consistent with *tub1*-R403C/H mutants disrupting dynein-mediated spindle positioning. Importantly, the phenotypes of *tub1*-R403C/H mutants in this assay are less severe than complete loss of dynein, as combining *dyn1*Δ with *bim1*Δ led to only one viable double mutant with very slow growth (Table S5). In addition, we found that double mutants combining *tub1*-R403C or R403H mutations with *dyn1*Δ do not exhibit growth defects that are more severe than the individual mutants alone (Figure S3C and S3D). This confirms that R403C/H mutants disrupt, but do not completely ablate dynein function.

The disruption of dynein activity could be caused by a failure of dynein to be recruited to the microtubule plus-end. Dynein is recruited through interactions with binding partners, and independent of its own microtubule-binding activity (Carvalho et al. 2004; Lee et al. 2003; Markus et al. 2009). We assessed dynein recruitment by measuring the intensity of GFP-tagged dynein on the plus-ends of microtubules (Figure 4E and 4F). We find that dynein localization is not altered in the tub1-R403C/H cells, and that the GFP signal intensity is not significantly different in either mutant condition (R403C p=0.96, R403H p=0.38, by t-test; Figure 4F). Therefore, the dynein defect observed in *tub1*-R403H and R403C mutants is not likely to be caused by a failure to recruit dynein to microtubule plus-ends.

Dynein and kinesin bind to nearby regions of the microtubule surface (Mizuno et al. 2004). We therefore asked whether *tub1*-R403C/H mutants also disrupt the activities of kinesins. We measured the intensity of the kinesin-8 motor Kip3 labeled with tdTomato in *tub1*-R403C/H yeast strains (Figure 4G and 4H). We observed no significant difference in the intensity of Kip3-tdTomato on microtubule plus-ends in either mutant condition compared to wild type, indicating that Kip3 binding and plus-end motility are not altered by tub1-R403C/H (p=0.8972 for R403C, p=0.8576 for R403H, by t-test; Figure 4H). This suggests that R403C and H substitutions disrupt dynein specifically and do not lead to wide-spread interference with interactions between microtubules and all motors.

### Dynein activity scales with amount of Tub1-R403H present in cells

Finally, we asked whether the dynein defect scales with the abundance of mutant α-tubulin in microtubules. Our results show that dynein activity is significantly disrupted in *tub1*-R403C/H haploid cells that also express the wild-type *TUB3* α-tubulin isotype (Figure 4A-4D and S5C-S5F). We sought to analyze dynein activity in cells expressing only mutant α-tubulin, by generating haploid cells that contain tub1-R403H α-tubulin but lack the minor α-tubulin isotype *TUB3* (*tub1*-R403H *tub3*Δ). *tub1*-R403H is viable as the only copy of α-tubulin, but *tub1*-R403C mutant cells do not survive in the absence of *TUB3* (Table S5). Microtubules in *tub1*-R403H *tub3Δ* mutant cells collide with the cortex less frequently than in *tub1*-R403H with a wild-type *TUB3* (Figure 5A). Even taking this into consideration, the tub1-R403H *tub3Δ* mutant cells exhibit a more severe loss of dynein phenotype than cells expressing *tub1*-R403H in combination with wild-type *TUB3* (Figure 5B). This indicates that the presence of some wild-type α-tubulin partially supports dynein function. As a second approach, we assayed diploid yeast strains that were either homozygous for wild-type *TUB1*, heterozygous for *tub1*-R403H, or homozygous for *tub1*-R403H. In this experiment, all the genotypes analyzed include two copies of *TUB3*. Of these diploid strains, the homozygous tub1-R403H mutant exhibits the most severe disruption of dynein activity (Figure 5C). The heterozygous mutant strain shows a slight, but not significant, defect in dynein activity compared to wild-type. Therefore, our results demonstrate that dynein disruption by *tub1*-R403H scales with the relative abundance of mutant α-tubulin in the cell.

**Figure 5.**
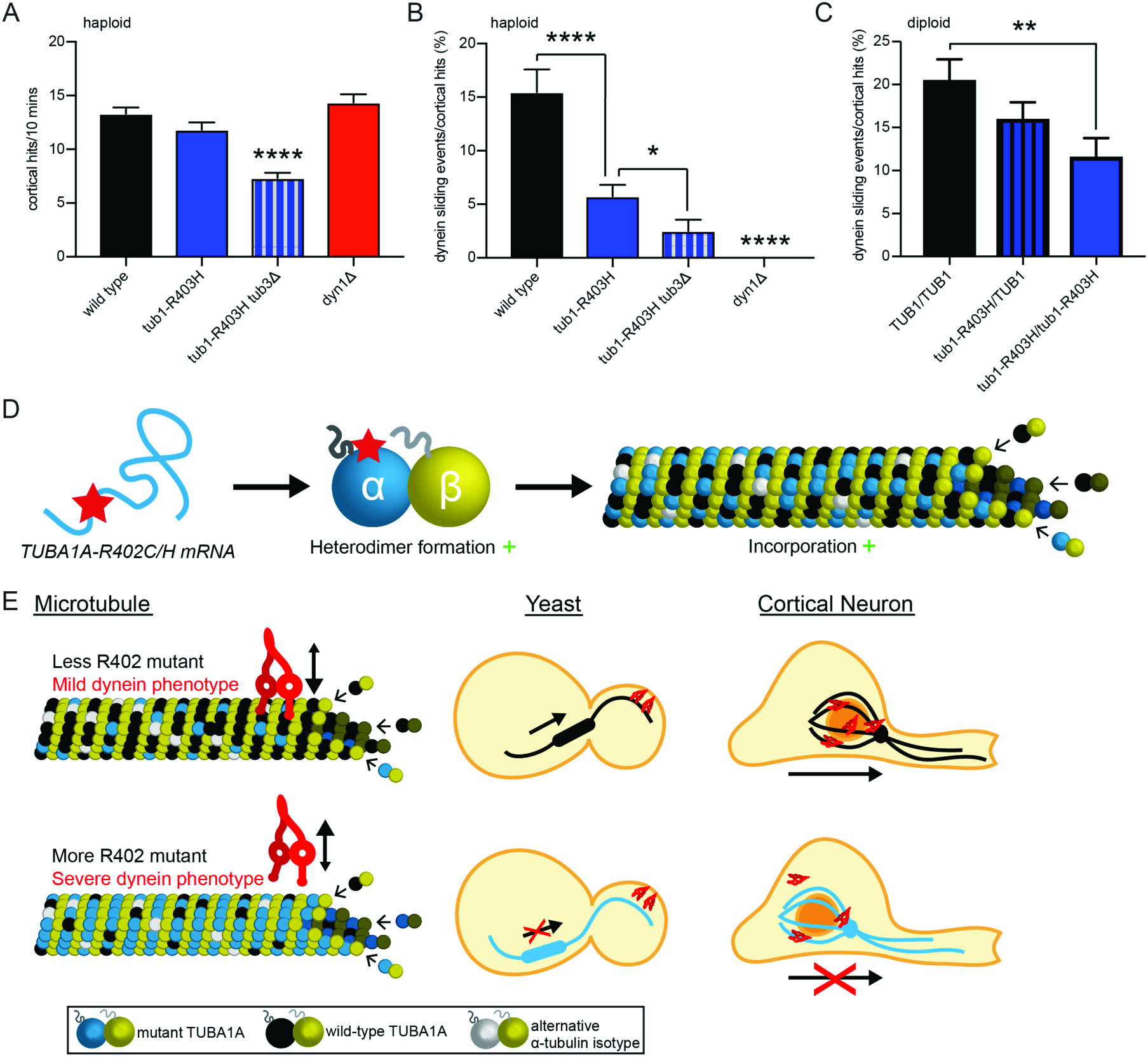
Dynein activity disruption scales with abundance of α-tubulin R402 mutant. (A) Quantification of microtubule-cortical hits in a 10-minute period for the indicated strains. (B) Quantification of dynein sliding events normalized to microtubule-cortical hits in haploid cells with varying levels of *tub1*-R403H mutant, displayed as percentage. At least 27 cells were analyzed for each strain. Data are represented as mean ± SEM. Single asterisks indicate significant difference compared to wild type, by *t*-test (*p*<0.05). Quadruple asterisks indicate significant difference compared to wild type, by *t*-test (*p*<0.0001). Strains: wild type, yJM1887, yJM1237; tub1-R402H, yJM2218; tub1-R402H tub3Δ, yJM2654; dyn1Δ, 1023. (C) Quantification of dynein sliding events normalized to microtubule-cortical hits in diploid cells with varying levels of tub1-R403H mutant, displayed as percentage. At least 30 cells were analyzed for each strain. Data are represented as mean ± SEM. Strains: wild type, yJM2711; tub1-R402H/TUB1, yJM2655, yJM2656; tub1-R402H/tub1-R402H, yJM2712. (D-F) Model of α-tubulin R402C and R402H mechanism of action. (D) R402C/H mutants form appropriate tubulin heterodimers that incorporate into microtubules. (E) Disruption of dynein:microtubule interaction scales with abundance of R402C/H mutant in the microtubule. With less R402C/H mutant incorporation, dynein actively slides microtubules to orient the mitotic spindle in yeast cells and neurons can migrate in the developing cortex. With more R402C/H mutant incorporation, dynein cannot appropriately orient the mitotic spindle in yeast cells and neurons fail to migrate out of the ventricular zone.

## DISCUSSION

Our finding that ectopic expression of *TUBA1A*-R402C/H alleles in the developing mouse cortex causes neuronal migration defects strongly supports a causal role of human *TUBA1A* mutations in lissencephaly. We provide insights into how *TUBA1A* substitutions at conserved residue R402 impact molecular level interactions with dynein. Furthermore, our results indicate that TUBA1A-R402C/H act dominantly to disrupt microtubule function, rather than leading to haploinsufficiency as predicted for other previously described *TUBA1A* mutations identified in mouse models (Keays et al. 2007; Belvindrah et al. 2017; Gartz Hanson et al. 2016). Based on the marked defect in cortical migration (Figure 1 and S1) and the inhibition of dynein activity by tub1-R403 mutant yeast cells (Figure 4A-4D and S5C-S5F), we propose a model in which *TUBA1A*-R402C/H mutants are assembled into microtubules and ‘poison’ the lattice by failing to support dynein activity, and this dynein impairment ultimately leads to migration failure during brain development (Figure 5D and 5E).

The molecular consequences of many of the patient-derived *TUBA1A* mutations are unknown. Previous data has provided evidence that many identified *TUBA1A* mutations may lead to haploinsufficiency, resulting in fewer stable tubulin heterodimers for neurons to use for important microtubule functions (Tian et al. 2010; Kumar et al. 2010). Indeed, two previous studies provided insight into the capability of TUBA1A-R402C/H to form functional tubulin heterodimers. *In vitro* transcription/translation analysis in rabbit reticulocyte to test tubulin heterodimer yield suggested that *TUBA1A*-R402C mutants limit the pool of functional heterodimers, while *TUBA1A*-R402H only leads to a slight reduction in tubulin heterodimer (Tian et al. 2010). However, despite destabilizing heterodimers *in vitro*, when expressed in cells mutant TUBA1A-R402C is capable of incorporating into microtubules in P19 cells (Kumar et al. 2010). Thus, whether R402 mutants in patients deplete the pool of functional heterodimers, or the mutant heterodimers assemble into microtubules and disrupt important microtubule functions of the microtubule lattice is a point of controversy. Our findings point to an alternative model – that *TUBA1A*-R402C and -R402H mutants act in a dominant fashion, forming tubulin heterodimers that incorporate into microtubules and disrupting neuronal migration even when appropriate levels of wild-type *TUBA1A* are present (Figure 5D and 5E).

Our findings reveal that *TUBA1A*-R402C/H mutants are capable of acting dominantly, appropriately incorporating into microtubules and disrupting dynein function. It is important to note that in our neuronal migration studies, cortical neurons contain two endogenous copies of wild-type *Tuba1a*. If the *TUBA1A*-R402C/H alleles identified in patients acted only to diminish the amount of functional tubulin heterodimer within a cell, then ectopic expression of mutant alleles would not be predicted to alter cortical formation in this system. On the contrary, we find that ectopic expression of *TUBA1A*-R402C/H mutants dominantly disrupts neuron migration, causing neurons with high mutant vector expression to be stuck in the ventricular zone. Interestingly, we observed that neurons ectopically expressing *TUBA1A*-R402C/H at a low level, as indicated by cytosolic GFP fluorescence intensity, are able to migrate to the cortical plate (Figure 1E-1G and S1B). This suggests that the neurons can overcome the dynein-inhibitory effect of the *TUBA1A*-R402C/H mutants if they contain a high enough ratio of wild-type tuba1a:TUBA1A-R402C/H protein. Interestingly, the *TUBA1A*-R402H neurons that reached the cortical plate exhibited significantly less GFP fluorescence than *TUBA1A*-R402C expressing neurons in the cortical plate, suggesting that *TUBA1A*-R402H may be more detrimental to neuronal migration than *TUBA1A*-R402C. These data are consistent with published patient data in which patients with *TUBA1A*-R402H mutations tend to display more severe lissencephaly phenotypes than patients with *TUBA1A*-R402C mutations (Poirier et al. 2007; Fallet-Bianco et al. 2008; Morris-Rosendahl et al. 2008; Kumar et al. 2010; Bahi-Buisson et al. 2014; Kamiya et al. 2014; Keays et al. 2007). In budding yeast, the interference of dynein activity also scales with the percentage of mutant α-tubulin in the cell. Increasing the relative abundance of *tub1*-R403H α-tubulin in the cell leads to more severe reduction of dynein sliding events (Figure 5B and 5C). Dynein-mediated sliding activity is diminished in haploid *tub1*-R403C/H cells, and decreases even further in haploid *tub1*-R403H mutant cells lacking the alternative α-tubulin isotype *TUB3*. Similarly, in diploid cells, the heterozygous mutant *tub1*-R403H/TUB1 trends towards decreased dynein sliding frequency, but the decrease does not become significant until both *TUB1* genes harbor the R403H substitution. Together, these data predict that the tubulin composition in the developing brains of patients, especially the levels of mutant *TUBA1A* compared to wild-type α-tubulin, can contribute greatly to the severity of brain malformation observed.

Why might these specific molecular perturbations to *TUBA1A* be uniquely detrimental to cortical development? Expression of α-tubulin isotypes is tightly regulated both by cell type and through developmental time. *TUBA1A* is highly upregulated in post-mitotic neurons during development, but is not found in neuronal progenitors, glia cells, or other organ systems, excepting one report of low *TUBA1A* mRNA levels in the testes and lungs (Lewis et al. 1985; Miller et al. 1987; Gloster et al. 1994; Coksaygan et al. 2006; Gloster et al. 1999; Aiken et al. 2017). The tight restriction of *TUBA1A* expression to developing post-mitotic neurons accounts for the human phenotypes being limited to the nervous system, with cortical malformation the hallmark of *TUBA1A* disorders. *TUBA1A* is not the only tubulin isotype to have been linked to tubulinopathies, as mutations have been identified in β-tubulin isotypes *TUBB2B*, *TUBB3*, and *TUBB5*, but *TUBA1A* is the only α-tubulin isotype found to associate with human disorders. This suggests that either 1) mutations to other α-tubulin isotypes may not lead to dominant phenotypes during brain development, or 2) mutations to other, more ubiquitously expressed α-tubulin isotypes may disrupt development so severely, perhaps by perturbing mitotic division, that they are not compatible with life. The restriction of disease-causing mutations to *TUBA1A* points to this isotype being particularly important for neurodevelopment.

Given the strong disruption of *TUBA1A*-R402C/H mutants on cortical migration, we were surprised to find only weak phenotypes in primary neuronal culture, despite using the same expression vector scheme for both sets of experiments. We noted a small, but significant increase in the ratio of microtubules oriented with the polymerizing plus-end toward the soma in the axons of neurons expressing TUBA1A-R402C or TUBA1A-R402H compared to in the axons of neurons expressing TUBA1A-WT or GFP-MACF43 alone. Orientation of plus ends away from the soma is partially dependent on dynein (del Castillo et al. 2015; Arthur et al. 2015). We determined that neurite formation, axon specification, axonal trafficking, and axonal microtubule density and polymerization were not significantly altered in primary cortical neurons ectopically expressing *TUBA1A*-R402C/H alleles (Figure 2, S2, and S3). Surprisingly, we also found that *TUBA1A*-P263T, a previously interrogated mutant which has been shown to form stable heterodimers, incorporate into microtubules, and disrupt polymerization (Tian et al. 2010), also did not decrease microtubule polymerization rates in primary cortical axons (Figure 2B and 2C). Our experiments were designed to minimize potential sources of variability between conditions. In our system, we express untagged *TUBA1A* alleles and fluorescently tagged MACF43 from the same vector. We selected GFP-MACF43 to mark the growing plus-ends of microtubules instead of introducing fluorescently tagged EB proteins. While the addition of ectopically expressed EB1 increases microtubule polymerization rate (Vitre et al. 2008), MACF43 acts as a more inert marker by interacting with endogenous EB1 to track microtubule tips (Honnappa et al. 2009). Thus, our microtubule polymerization rates are slower than previous reports utilizing an ectopic *TUBA1A* expression system with tagged EB proteins (Tian et al. 2010; Belvindrah et al. 2017), and are consistent with rates observed in mammalian neuronal systems using GFP-MACF43 (Yau et al. 2016). We carefully controlled for the neuronal environment that we sampled, including only rates of anterograde microtubules polymerizing in DIV11 axons a consistent distance from the soma. Importantly, we conducted multiple sets of experiments encompassing numerous primary neuron preparations and vector nucleofection. Over the course of the experiments, we observed a large spread in microtubule polymerization rate data (Figure 2C). We observed cells with slower polymerization and cells with faster polymerization in each condition, with cells from the same neuronal preparation exhibiting similar rates. This suggests that there is inherent variability between primary neuron preparations that must be taken into consideration when analyzing microtubule polymerization events. Based on these data, we conclude that even when evaluating mutants that should form stable heterodimers and polymerize into microtubules (Figure 3D-3F; Tian et al., 2010), ectopic expression of *TUBA1A* mutant alleles may not be sufficient to disrupt the axonal microtubule network in primary cortical cultures.

The drastic disruption to cortical migration by ectopic expression of *TUBA1A*-R402C/H mutant alleles, but not to primary cortical neuron function, suggests that newly-born, migrating neurons are uniquely sensitive to *TUBA1A* perturbation. This sensitivity may be attributable to the importance of dynein in neuronal migration, which is not observed in cultured neurons. Alternatively, there may be differences in the levels of mutant *TUBA1A*-R402C/H relative to wild-type *Tuba1a* mutant in these systems, with a higher ratio of wild-type *Tuba1a* able to mask the consequences of the *TUBA1A*-R402C/H mutants in the primary neurons. Consistent with this, our findings indicate that levels of mutant tubulin may determine the severity of the phenotype (Figure 1E-1G, 5B, and 5C). Tubulinopathy patients express heterozygous *TUBA1A* mutations; therefore, approximately half of the available *TUBA1A* in patients is mutant *TUBA1A*. In contrast, the neurons analyzed in this study express exogenous *TUBA1A* mutants from a plasmid in addition to two endogenous wild-type *TUBA1A* copies present. It is therefore important for researchers to consider novel methods to study tubulin mutants in a system containing endogenous mutant tubulin expression.

Mutations that alter the same residue of TUBA1A but lead to different side-chain substitutions have been identified in tubulinopathy patients, but the molecular consequence of the different substitutions remains unclear. While both the spectrum of lissencephaly phenotypes observed in patients and the cortical migration phenotypes caused by the two substitutions, *TUBA1A*-R402C and *TUBA1A*-R402H, are nearly indistinguishable (Figure 1 and 4), we find that replacing the native arginine with cysteine or histidine lead to distinguishable molecular consequences for α-tubulin function. Whereas replacing the native arginine with histidine, another basic amino acid, mainly leads to dynein disruption with no other obvious phenotypes seen in yeast (Figure 4), replacing arginine with cysteine, a neutral, nucleophilic residue, not only disrupts dynein but also leads to additional phenotypes in yeast growth and genetic assays (Table S5 and Figure S5). *tub1*-R403C, but not *tub1*-R403H, leads to a drastic reduction of yeast growth when cells are stressed with the microtubule destabilizing drug benomyl, and is not viable as the sole copy of α-tubulin (Figure 5A-C, 5E, and Table S5). This difference may be due to the positively-charged arginine providing important intra-tubulin stabilization by forming a salt bridge with E416 on the negatively-charged surface of α-tubulin, which has been proposed based on structural data (Uchimura et al. 2015). The neutral cysteine substitution may disrupt this salt bridge while the basic histidine supports it. This suggests that tubulinopathy phenotypes arise not merely from the loss of the native residue, but also from the gain of the substituted side-chain. This is important when considering several other patient-derived *TUBA1A* mutants that affect the same residues but lead to the introduction of different side chains.

In conclusion, patient mutations identified in *TUBA1A* at residue R402 disrupt dynein activity and dominantly cause a pronounced cortical neuron migration defect. Our results demonstrate that the *TUBA1A-R402* mutants are causal for the lissencephaly phenotype observed in patients, and that the amount of *TUBA1A*-R402C/H in the microtubule network contributes to the severity of the phenotype. This work reveals that patient-derived mutations to tubulin genes can provide important insight into the molecular interactions with microtubules to enable correct brain development. Future studies into additional mutations may shed light on other important microtubule-mediated functions in neurodevelopment.

## MATERIALS AND METHODS

### Neuronal Expression Vectors

Expression constructs used in this study are listed in Table S7. The coding region of human *TUBA1A* from the Mammalian Genome Collection (clone ID:LIFESEQ6302156) was cloned into the multiple cloning site of pCIG2 plasmids using sticky-end cloning with XhoI and PstI. QuikChange Lightning Site-Directed Mutagenesis Kit (Agilent) was used to introduce the p.R402C or p.R402H substitution to *TUBA1A*. Mutations were confirmed by sequencing across the complete *TUBA1A* coding region. For experiments measuring microtubule polymerization in cells expressing *TUBA1A* mutants, GFP-MACF43 was cloned into the pCIG2-*TUBA1A* constructs. GFP-MACF43 construct was generously gifted by Casper Hoogenraad and Laura Ann Lowry, and we used these to create GFP-MACF43-expressing pCIG2 plasmids using sticky-end cloning to place MACF43 after the encoded GFP with BsrGI and NotI. To assess axonal trafficking, a mNeonGreen-Lysosomes-20 (Rattus norvegicus Lysosomal Membrane Glycoprotein 1 (LAMP1; NM_012857.1)) fusion was obtained from Allele Biotechnology (Shaner et al Nat Methods 2013), and cloned into pCIG2 using NEB Gibson Assemby Cloning Kit (NEB #E2611) to replace cytosolic GFP. Oligos used for plasmid construction are listed in Table S8. The same pCIG2 constructs were used for both the *in vivo* migration assay and the primary neuronal culture morphology assay, with neurons producing cytosolic GFP and mutant, wild-type, or no additional *TUBA1A* under the same expression regulation.

### *In utero* electroporation

*In utero* electroporation was performed on E14.5 embryos from timed pregnant mice. Endotoxin-free plasmid DNA (see ‘Neuronal Expression Vectors’ section, above) was prepared with fast green dye (1% stock) to a concentration of 1 μg/μl in TE buffer (10mM Tris Base, 1mM EDTA Solution, pH 8.0). Plasmid DNA was injected into the lateral ventricles of the exposed embryos, and electroporated with 5 pulses at 40 V separated by 900 ms. Embryos were returned and allowed to develop to E18.5, at which point their brains were dissected and fixed in 4% paraformaldehyde overnight at 4°C. Brains were sectioned coronally with a vibrating microtome (VT1200S; Leica) at 50 um thick sections. Sections were then mounted, imaged on a Nikon spinning disc confocal microscope with a 0.3 NA 10x Plan Fluor objective, and analyzed to assess migration of electroporated cells. Analysis includes at least 4 animals for each condition.

### Yeast strains and manipulation

General yeast manipulation, media, and transformation were performed by standard methods (Amberg et al., 2005). Strains are provided in Table S6. *TUB1*-R403C and *TUB1*-R403H mutant alleles were generated at the endogenous *TUB1* locus using methods described by Toulmay and Schneiter (2006). The selectable marker *URA3* was inserted at the *TUB1* locus following the 3’UTR. An amplicon including *TUB1*, 3’UTR, and URA3 marker was then amplified from genomic DNA by PCR, using mutagenic oligos to introduce R403C or R403H mutations, and transformed into a wild-type strain background. Mutations were confirmed by sequencing over the complete *TUB1* genomic locus. GFP-Tub1 fusions were under the *TUB1* promoter and integrated at the *LEU2* locus and expressed ectopically, in addition to the native, untagged *TUB1* (Song and Lee, 2001). Kip3-tdTomato, Kip2-mEmerald, and Bim1-mNeonGreen fusion were generated by PCR-mediated tagging at the native loci (Sheff and Thorn, 2004). The mNeonGreen plasmid DNA was originally acquired from Allele Biotechnology and Pharmaceuticals (Shaner et al. 2013). Dyn1-3GFP was generated at the native *DYN1* locus using an integrating plasmid (Lee 2003 JCB). *TUB3* was deleted by replacing the native coding sequence with the KanMX6 marker, using conventional PCR-mediated methods (Petracek & Longtine 2002). *DYN1* and *BIM1* were knocked out using the same method, but replaced with HIS3MX6. Oligos used for plasmid and yeast strain construction are listed in Supplemental Table 8.

### Yeast growth assays

Cells were grown to saturation at 30°C in 4 ml YPD (Yeast extract, Peptone, Dextrose; rich media) and then a 10-fold dilution series of each strain was spotted to a YPD plate and a YPD plate supplemented with 10 μg/ml benomyl (Sigma, #381586). YPD plates were incubated at 30°C for 3 days, and benomyl supplemented plates were grown at 30°C for 4 days.

### Western blots of α-tubulin levels in yeast

Cell cultures were grown shaking at 30°C until log phase in 5 ml YPD. Cells were then resuspended in chilled lysis buffer (6mM Na_2_HPO4, 4mM NaH_2_PO_4_, 1% NP40, 150mM NaCl, 2mM EDTA, 50mM NaF, 4μg/μL Leupeptin, 0.1mM Na_3_VO_4_) with 1X fungal protease inhibitor cocktail (Sigma, #P8215), and lysed by 5 cycles of bead beating at 4°C. Cells were centrifuged to obtain the clarified lysate. Total protein concentration was determined by Bradford assay, and all samples were normalized to 1.2 μg/μL. Samples were prepared with Laemmli sample buffer and β-mercaptoethanol, and then 24μg of each lysate was loaded per lane onto an 10% SDS PAGE gel. After running the gels, the samples were transferred to a PVDF membrane and blocked in Odyssey (LI-COR; product # 927-40000) buffer overnight at 4°C. Membranes were probed with mouse-anti-α-tubulin (4A1; AB_2732839; 1:100; (Piperno and Fuller, 1985)) and rabbit-anti-Zwf1 (Sigma A9521; <u>AB_258454</u>;1:10,000), followed by secondary antibodies goat-anti-mouse-680 (LI-COR 926-68070, Superior, NE; 1:15,000) and goat-anti-rabbit-800 (LI-COR 926-32211; 1:15,000), and imaged on an Odyssey Imager (LI-COR Biosciences). α-tubulin band intensity was analyzed in ImageJ and the intensity of the slower migrating Tub1 isotype was compared to the amount of total α-tubulin.

### Yeast microscopy and image analysis

For time-lapse imaging of microtubule dynamics and sliding, living cells were grown asynchronously at 30°C to early log phase in nonfluorescent medium, mounted on coverslips coated with concanavalin A (2 mg/mL; Sigma #C2010), and sealed with VALAP (vasoline:lanolin:paraffin at 1:1:1) (Fees et al. 2017). Imaging was performed on a Nikon Ti-E microscope equipped with a 1.45 NA 100X CFI Plan Apo objective, piezo electric stage (Physik Instrumente, Auburn, MA), spinning disc confocal scanner unit (CSU10; Yokogawa), 405-, 488-, 561-, and 640-nm lasers (Agilent Technologies, Santa Clara, CA), and an EMCCD camera (iXon Ultra 897; Andor Technology, Belfast, UK) using NIS Elements software (Nikon) (hereafter referred to as Nikon spinning disc confocal microscope). During acquisition, stage temperature was 25°C. Images were analyzed using ImageJ (National Institutes of Health, downloaded from https://imagej.nih.gov).

### Yeast microtubule dynamics analysis

Microtubule dynamics were analyzed as described (Fees et al. 2017). The lengths of astral microtubules were determined by the localization of the plus-end tracking protein Bim1-mNeonGreen, imaged at 4- or 5-s intervals for 10 minutes. Only astral microtubules from preanaphase cells were measured for this analysis. Polymerization and depolymerization events were defined as at least three contiguous data points that produced a length change ≥0.5 μm with a coefficient of determination ≥0.8. Microtubule dynamicity was calculated by the total change in length divided by the change in time and expressed in tubulin subunits changed per second (Toso et al., 1993). At least 15 astral microtubules were analyzed for each genotype. Dynamics measurements for individual microtubules were pooled by genotype and then compared with pooled data for different genotypes using a Student’s t test to assess whether the mean values for different datasets were significantly different.

### Analysis of microtubule–cortex interactions and microtubule sliding

Cells expressing an integrated fusion of GFP to the N-terminus of α-tubulin (GFP-Tub1) were HU-treated, imaged and analyzed as described previously (Estrem et al, 2017). Briefly, cells were arrested with HU for 2 hours and imaged in time-lapse Z-series for 10 minute with 4-s time interval between frames. Microtubule-cortex interactions were identified in max-projections of Z-series images by the microtubule touching the cell cortex, which was identified by signal of free GFP-Tub1 subunits in the cytoplasm. Dynein sliding events were defined as spindle translocation coinciding with microtubule-cortex interaction, and the total distance and time of the spindle movement was measured.

### Localization of dynein, kinesin-8/Kip3

Two-color images of cells expressing Dyn1-3GFP and Spc110-DsRed, or Kip3-tdTomato and Spc110-GFP were collected in time-lapse, full-cell Z series separated by 0.4 um. Only preanaphase cells were used for this analysis. Z series were collapsed into maximum intensity projections and analyzed in ImageJ. We identified the astral microtubule plus end as foci of Dyn1-3GFP or Kip3-tdTomato that were separate from the spindle pole bodies and rapidly changed position over time, since plus ends will move over time with dynamic microtubules. In contrast, cortical foci of Dyn1-3GFP were stationary in time-lapse imaging. Intensities of Dyn1-3GFP or Kip3-tdTomato signals were measured within a 397 nm^2^ region (630 × 630 nm). These values were adjusted for background signal by taking a background fluorescence intensity measurement of the same size adjacent to the plus end of the microtubule and then subtracting that value from the value measured at the plus end.

### Primary cortical cultures

For primary cortical neuronal cultures, the frontal cortex was dissected from postnatal day 0-2 male and female neonatal Sprague Dawley rats. Frontal cortex was dissociated using papain digestion. After washing away papain with MEM (Life Technologies, 11090-081), cells were gently dissociated first with a wide-bore fire polished pipet, then with a narrow-bore polished pipet. Plasmid DNA was introduced to 4×10^6^ neurons using AMAXA rat nucleofection kit (Lonza, VPG-1003), and then 500,000 AMAXA-treated neurons were plated per poly-D-lysine-coated glass bottom dish (Willco Wells, HBSt-3522) in MEM supplemented with 10% FBS (Sigma), 1% penicillin/streptomycin (Gibco, 15070-063), and 25 μM L-glutamine (Gibco, 25030081). After 2 hours, media was replaced with fresh, 10% FBS supplemented MEM. After 24 hours, media was replaced with Neurobasal-A (NBA, Life technologies, 10888-022) supplemented with 2% B27 (10 ml vial in 500 ml MEM, Gibco, 17504-044) and 10 μM mitotic inhibitors (uridine/fluoro deoxyuridine, Sigma, U3003/F0503). Neurons were maintained in a humidified incubator at 37°C with 5% CO_2_. Cells were then fed with supplemented NBA on DIV7. Neurons were imaged and analyzed on DIV11.

### Neuron immunocytochemistry

Primary cortical neuronal cultures were prepared as described above. On DIV11, neurons were washed twice with 1x Phosphate Buffered Saline (PBS), fixed with 4% paraformaldehyde for 10 minutes at room temperature. Cells were then washed three times with 1x PBS, for 5 minutes each. Permeabilization was performed with 0.1% Triton/PBS for 10 minutes at room temperature, agitated and then washed with PBS. The cells were blocked in 5% BSA/PBS for 30 minutes at room temperature, with agitation. Immunostaining was performed using antibodies to SMI-312 (Biolegends, # 837904) and MAP2 (Novus Biologicals, #NB300-213) at concentrations of 1 ug/ml and 1:2000, respectively, for 2 hours in a humidified chamber. After primary antibody staining, cells were washed 3x with PBS. Secondary antibody staining with Alexa Flour 568 Donkey anti-Mouse IgG H+L (Life technologies, #A10037) and Cy5 Goat anti-Chicken IgY H&L (Abcam, # ab97147) in 1%NGS/PBS was performed for 1 hour in a dark, humidified chamber. After washes, cells were mounted with vectashield (Vector Laboratories, #H-1200) and imaged on a Nikon spinning disc confocal microscope with a 40X oil objective.

### Time-lapse microscopy and analysis of neurons

Primary cortical neuronal cultures were prepared as described above, with relevant plasmid DNA introduced with AMAXA rat nucleofector kit (Lonza). At DIV11, live primary cultured neurons were treated with an antibody to external Neurofascin (extNF, Anti-Pan-Neurofascin (external), NeuroMab clone A12/18) as previously described (Dumitrescu et al., 2016) to label the axon initial segment. Briefly, cells were treated with 50:50 conditioned media: fresh NBA with 50 uM APV (Tocris) to protect against cell death (Hogins et al., 2011). Cells were then placed in primary antibody solution (extNF, 1:200) diluted in NBA-APV for 5 minutes at 37°C. After three washes in NBA-APV, cells were placed in secondary antibody solution (Donkey antimouse A568, 1:500, Life technologies #A10037) for 30 seconds at room temperature, followed by three washes with NBA-APV. ACSF imaging media supplemented with 2 mM CaCl_2_ and 1 mM MgCl_2_ was added to the neurons, which were imaged on a Nikon spinning disc confocal microscope with a 1.3 NA 40X CFI Plan Fluor oil objective. Cells were kept at 37°C during image acquisition. Axons were identified by presence of high extNF, and were imaged for 4 minutes at 2.25 second intervals (to track GFP-MACF43 comets) using a 0.4 μm Z-step to cover the depth of the imaged axon or 4 minutes at 0.32 second intervals at one Z-plane (to observe Lyso-20-mNeon trafficking). The max-projected (if applicable) time-lapse videos were analyzed in ImageJ using the kymograph feature. GFP-MACF43 polymerization rates and Lyso-20-mNeon trafficking rates were determined from the slope of the line on the kymographs.

## ACKNOWLEDGEMENTS

We are grateful to Santos Franco (University of Colorado School of Medicine) for aiding with *in utero* mouse cortical electroporations. We thank Matthew Kennedy and Mark del Acqua (University of Colorado School of Medicine) for providing P0-2 rat cortex. We thank Matthew Kennedy for gifting us the pCIG2 vector and Laura Anne Lowery (Boston College) and Casper Hoogenraad (Utrecht University) for sharing the GFP-MACF43 vector. This work was supported by the National Science Foundation Graduate Research Fellowship 1553798 to J.A., National Institutes of Health 5R01GM112893-04 to J.K.M., a pilot grant from the Department of Cell and Developmental Biology, University of Colorado Denver, Anschutz Medical Campus awarded to J.K.M and E.A.B.

## AUTHOR CONTRIBUTIONS

J.A., J.K.M., and E.A.B. conceived the project, designed the experiments, interpreted results, and wrote the manuscript. J.A. performed the experiments.

## SUPPLEMENTAL INFORMATION

**Figure S1.**
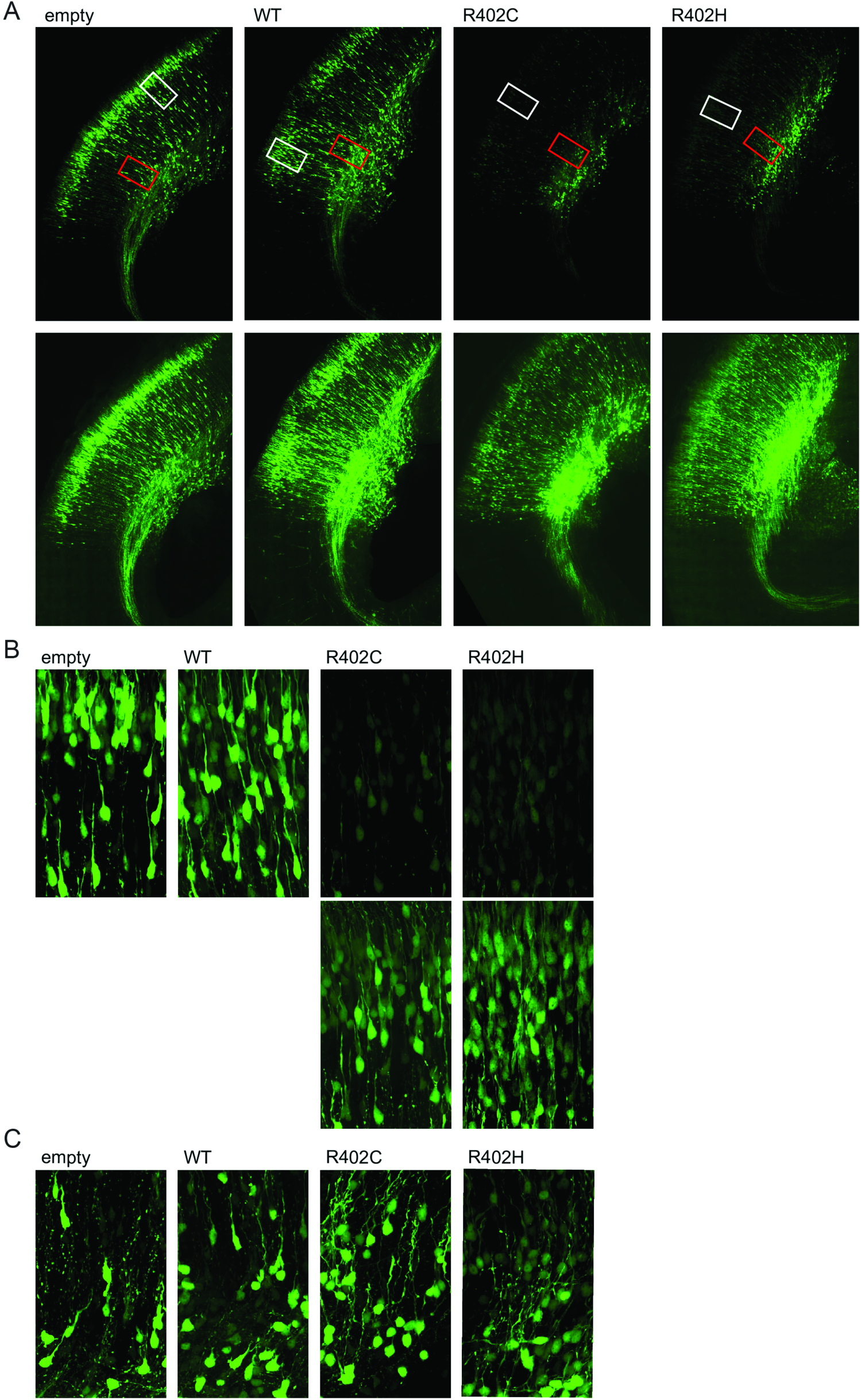
Neurons in the developing mouse cortex form neuronal projections when electroporated with empty vector and *TUBA1A* alleles. (A) Representative coronal sections with long axonal tracts from E18.5 mouse brain electroporated at E14.5 with pCIG2 vectors. Two exposures of the same sections are provided. (B) White insets from sections in (A) reveal cortical plate neurons with leading projections. Two exposures of the same R402C and R402H sections are provided. (C) Red insets from sections in (A) reveal cellular extensions in lower layer neurons.

**Figure S2.**
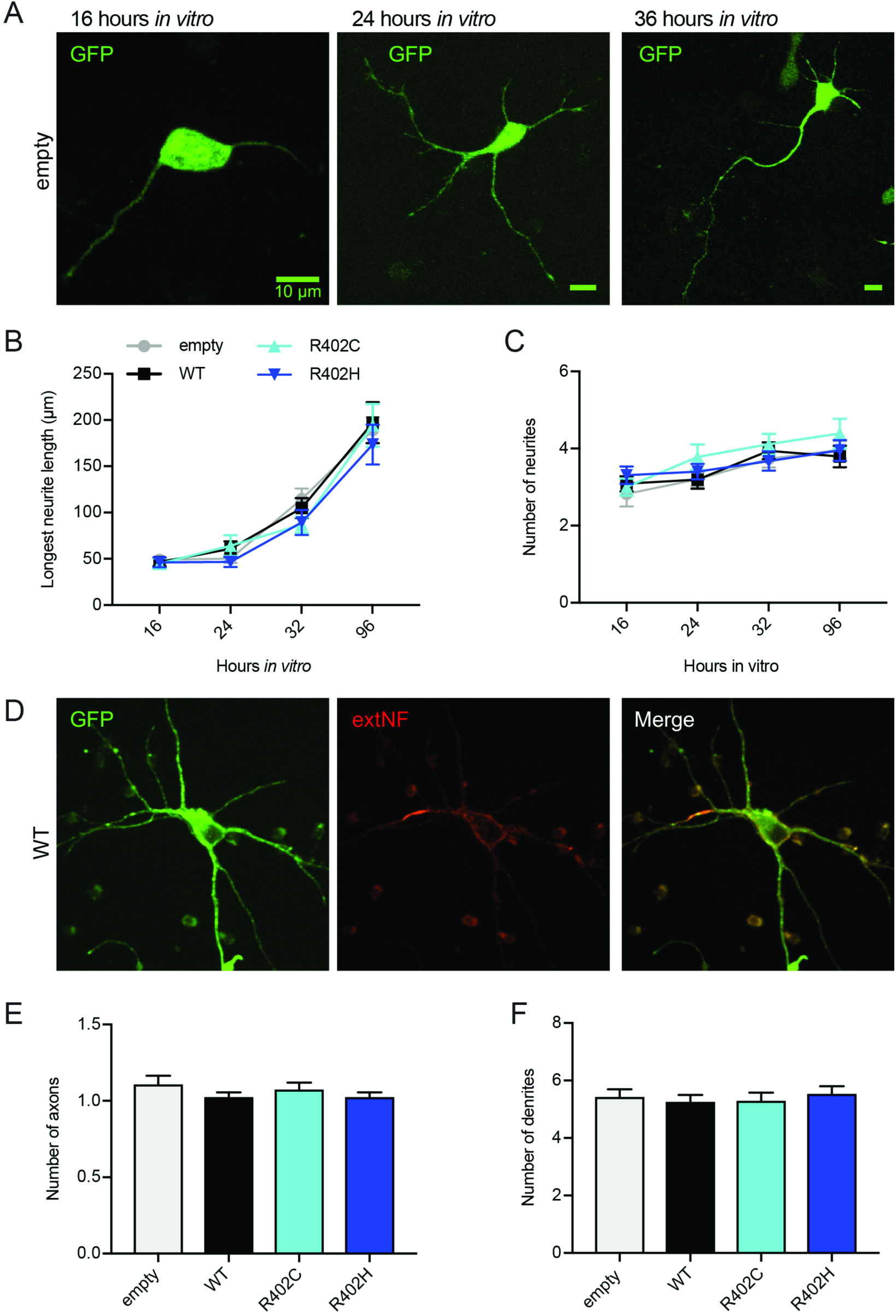
Neuron morphology is not significantly altered by ectopic expression of *TUBA1A*-R402C/H mutants. (A) Representative images of primary rat cortical neurons expressing pCIG2-*Tuba1a*(WT)-ires-GFP at 16 hours, 24 hours, and 36 hours *in vitro*. (B and C) Longest neurite length (B) or neurite number (C) tracked over 16 to 96 hours *in vitro* from neurons expressing pCIG2 vectors. (D) DIV11 neuron expressing pCIG2-Tuba1a(WT)-ires-GFP stained with external neurofascin (extNF) to mark the axon initial segment (AIS). (E and F) Number of axons (E) or dendrites (F) in neurons expressing pCIG2 vectors.

**Figure S3.**
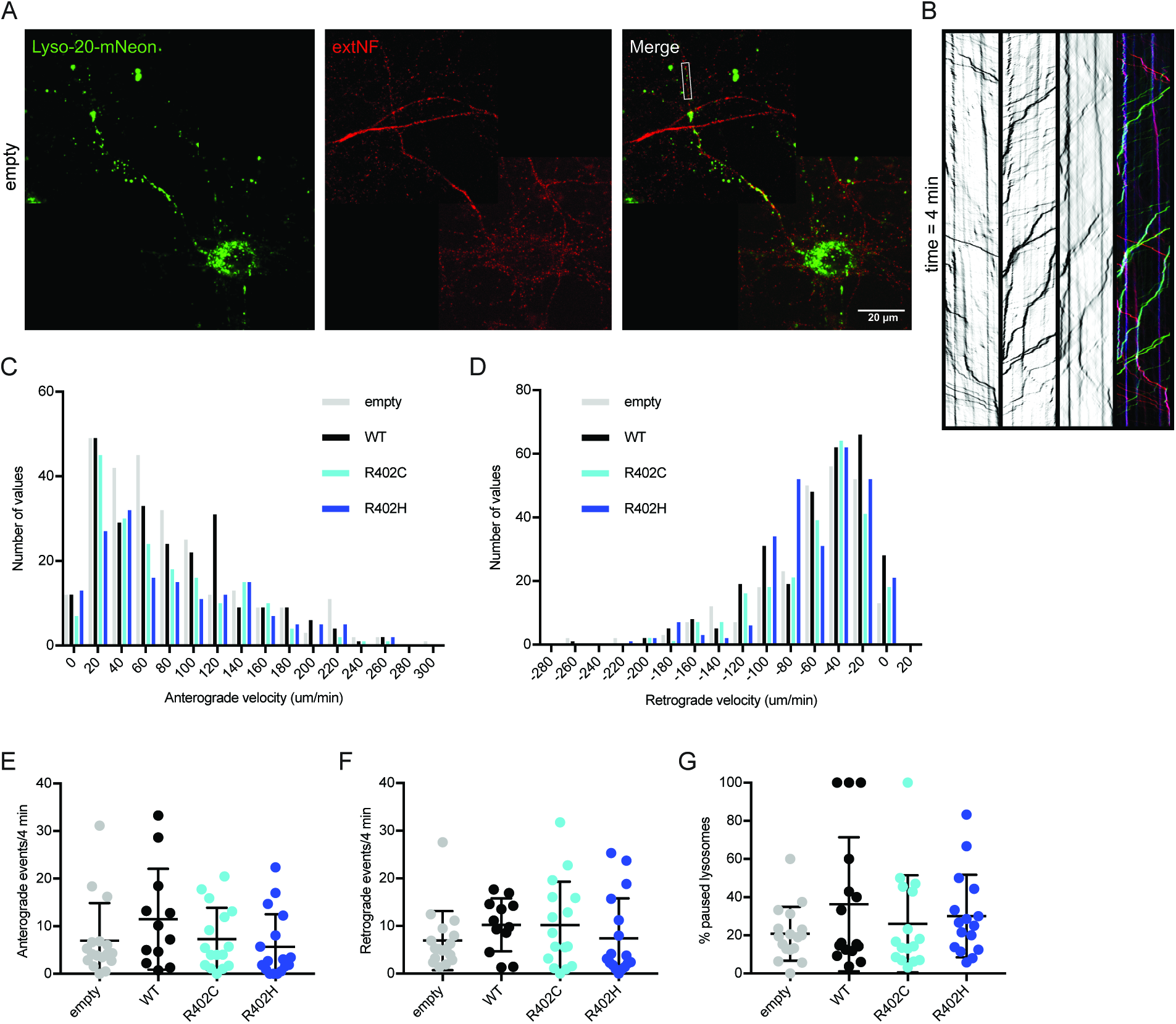
Axonal trafficking is not significantly altered by ectopic expression of *TUBA1A*-R402C/H mutants. (A) DIV11 primary rat cortical neuron expressing pCIG2-Lyso20-mNeon and live-stained with extNF to reveal AIS. (B) Kymograph of lysosome movement in inset from (A) showing anterograde (red), retrograde (green), and paused (blue) events over 4 minutes. (C and D) Histogram of anterograde (C) or retrograde (D) lysosome trafficking events observed in neurons expressing pCIG2-Lyso20-mNeon, pCIG2-Tuba1a(WT)-ires-Lyso20-mNeon, pCIG2-Tuba1a(R402C)-ires-Lyso20-mNeon, or pCIG2-Tuba1a(R402H)-ires-Lyso20-mNeon. (E-G) Number of observed anterograde (E), retrograde (F), or paused (G) lysosome trafficking events in 4 minutes.

**Figure S4.**
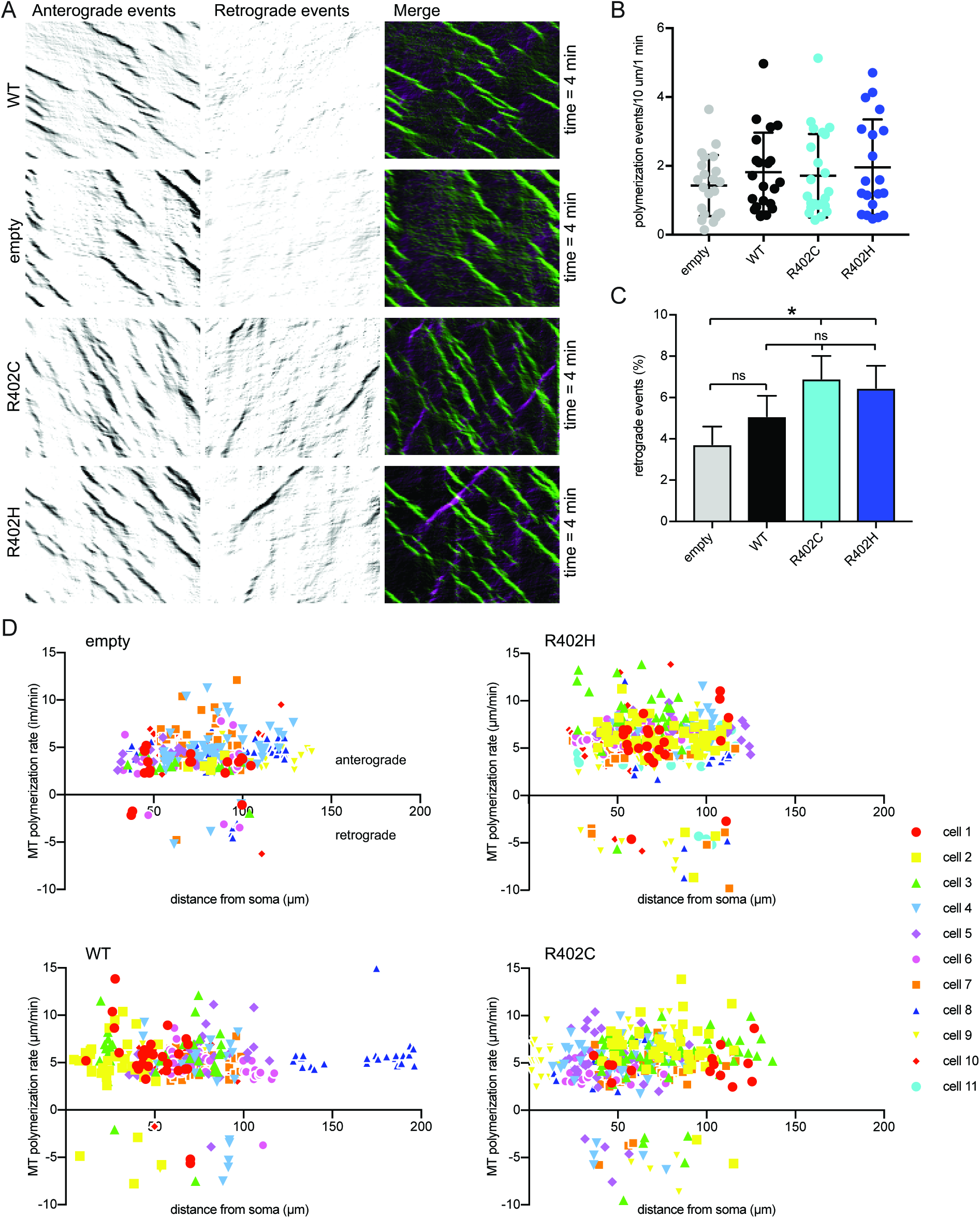
Axonal microtubule orientation may be altered by ectopic expression of *TUBA1A*-R402C/H mutants. (A) Representative kymographs revealing anterograde and retrograde axonal microtubule polymerization events in pCIG2-Tuba1a(WT)-ires-GFP-Macf43, pCIG2-GFP-Macf43, pCIG2-Tuba1a(R402C)-ires-GFP-Macf43, and pCIG2-Tuba1a(R402H)-ires-GFP-Macf43. (B) Number of polymerization events observed in neurons in a 10 μm region over 1 minute. Each data point represents cellular mean, with bars displaying mean ± SEM. No significant differences to empty vector or WT control, with significance determined as p<0.05. (C) Percentage of retrograde microtubule polymerization events. Data are represented as mean ± SEM. Asterisk indicates significance compared to empty vector control, as determined by Chi square analysis (p<0.05). (D) Plot of polymerization rates observed in axons by their distance from soma at initiation, with each data point representing an individual microtubule polymerization event. Anterograde rates are plotted as positive values, and retrograde rates as negative values.

**Figure S5.**
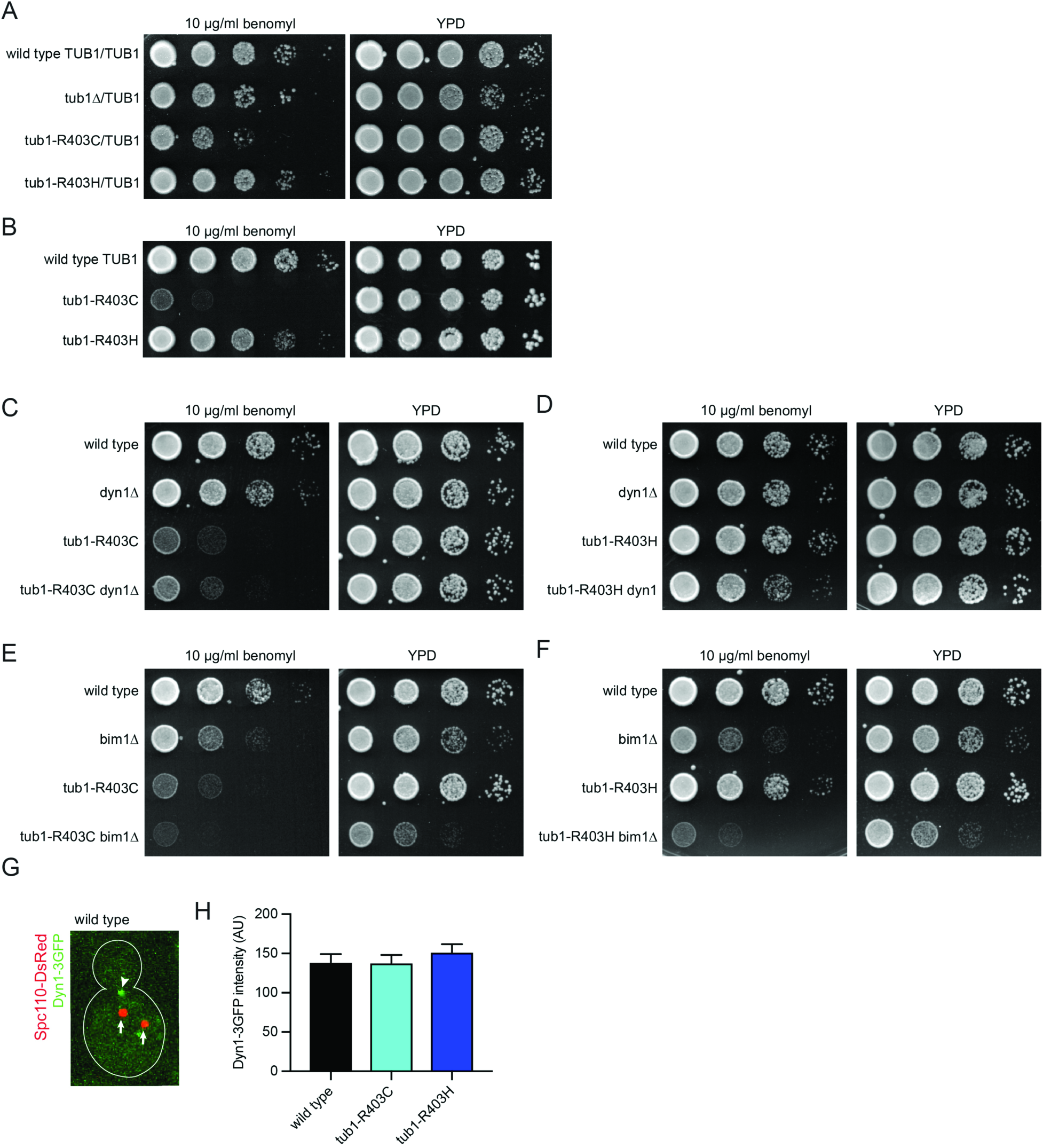
α-tubulin R402C/H mutants in *S. cerevisiae* support yeast growth and act in the dynein spindle positioning pathway. (A) Growth assay for heterozygous diploid *tub1-R403* mutants on rich media or rich media supplemented with the microtubule destabilizing drug benomyl (10 μg/mL), incubated at 30°C. Strains: wild type, yJM0091; tub1Δ/TUB1, 0591; tub1-R403C/TUB1, yJM2364, yJM2365; tub1-R403H, yJM2366, yJM2367. (B) Growth assay for haploid *tub1*-R403 mutants grown on rich media or benomyl supplemented rich media (10 μg/mL), incubated at 30°C. Strains: wild type, yJM1839, yJM1840; tub1-R403C, yJM2120, yJM2239; tub1-R403H, yJM2121, yJM2122. (C and D) Genetic interaction test of *tub1*-R403C (C) and *tub1*-R403H (D) mutants with *dyn1*Δ. Indicated strains were grown on rich media or rich media supplemented with benomyl (10 μg/mL), incubated at 30°C. Strains: wild type, yJM2256, yJM2261; dynM, yJM2255, yJM2264; tub1-R403C, yJM2258; tub1-R403C dyn1Δ, yJM2257; tub1-R403H, yJM2263; tub1-R403H dyn1Δ, yJM2262. (E and F) Genetic interaction test of *tub1*-R403C (E) and *tub1*-R403H (F) mutants with *bim1*Δ, a member of the compensatory spindle positioning pathway. Indicated strains were grown on rich media or rich media supplemented with benomyl (10 μg/mL), incubated at 30°C. Strains: wild type, yJM2243, yJM2252; bim1Δ, yJM2246, yJM2249; tub1-R403C, yJM2245; tub1-R403C bim1Δ, yJM2244; tub1-R403H, yJM2251; tub1-R403H bim1Δ, yJM2250.

**Table S1.** Axonal anterograde microtubule polymerization rates from DIV11 rat primary cortical neurons expressing *TUBA1A* vectors or controls. Data presented as mean ± SEM. 141-452 microtubule polymerization events were analyzed for each condition from at least 17 cells over a minimum of three separate neuron preparations. No significant differences to empty vector or WT control, with significance determined as *p*<0.05.

| | Microtubule polymerization rate (μm/min) |
| --- | --- |
| empty | 4.702±0.3212 |
| WT | 5.275±0.3371 |
| P263T | 5.801±0.2966 |
| R402C | 5.295±0.3033 |
| R402H | 5.469±0.2845 |

**Table S2.**
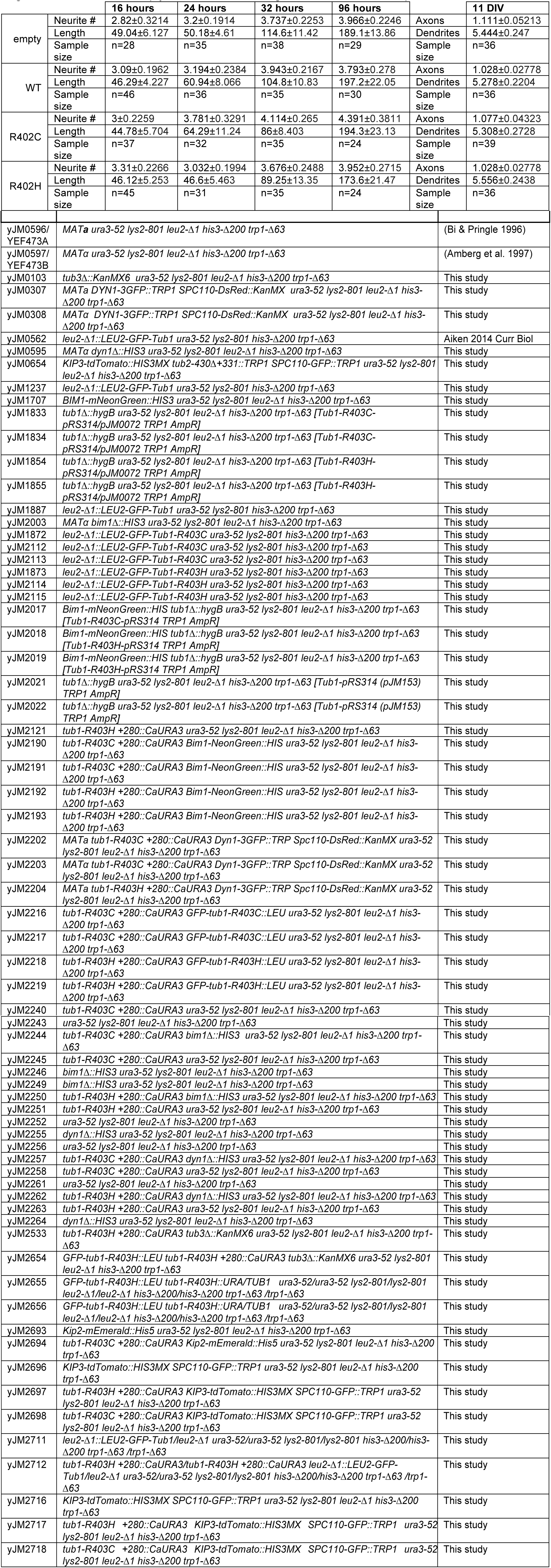
Neurite number and length over time *in vitro* from for rat primary cortical neurons expressing *TUBA1A* vectors or controls. Axons determined either by SMI312 staining or AIS marker extNF. Dendrites determined by MAP2 staining or absence of extNF staining. Data presented as mean ± SEM. No significant differences to empty vector control, with significance determined as *p*<0.05.

**Table S3.** Axonal lysosome trafficking events and rates from DIV11 rat primary cortical neurons expressing *TUBA1A* vectors or controls. Data presented as mean ± SEM. At least 16 cells were analyzed for each condition over three separate neuron preparations. No significant differences to empty vector control, with significance determined as *p*<0.05.

|  | Anterograde rate (μm/min) | Number anterograde events | Retrograde rate (μm/min) | Number retrograde events | Percent paused |
| --- | --- | --- | --- | --- | --- |
| empty | 78.55±3.749 | 6.952±1.92 | 61.97±2.985 | 6.945±1.501 | 20.82±3.42 |
| WT | 80.36±3.741 | 11.48±3.061 | 58.81±2.682 | 10.25±1.604 | 36.25±8.77 |
| R402C | 71.18±3.961 | 7.33±1.641 | 59.53±2.744 | 10.19±2.281 | 26.04±6.36 |
| R402H | 80.39±4.812 | 5.648±1.66 | 59.6±2.569 | 7.393±2.036 | 30.08±5.40 |

**Table S4.**
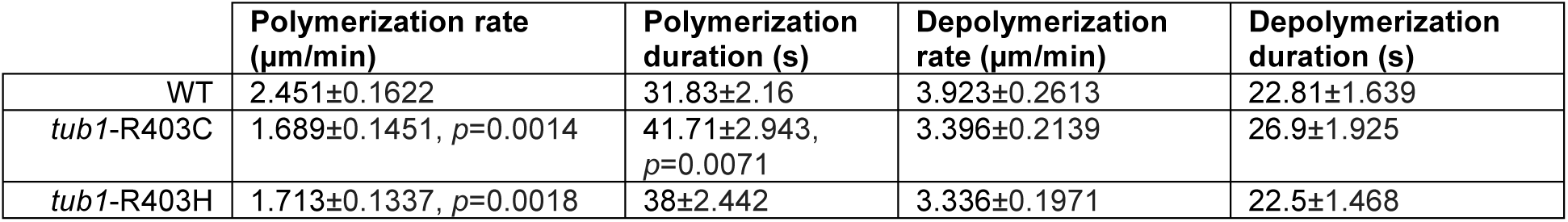
Yeast microtubule dynamics data. Data presented as mean ± SEM. At least 15 cells were analyzed for each condition. Unless p-value is provided, *p*>0.05.

**Table S5.**
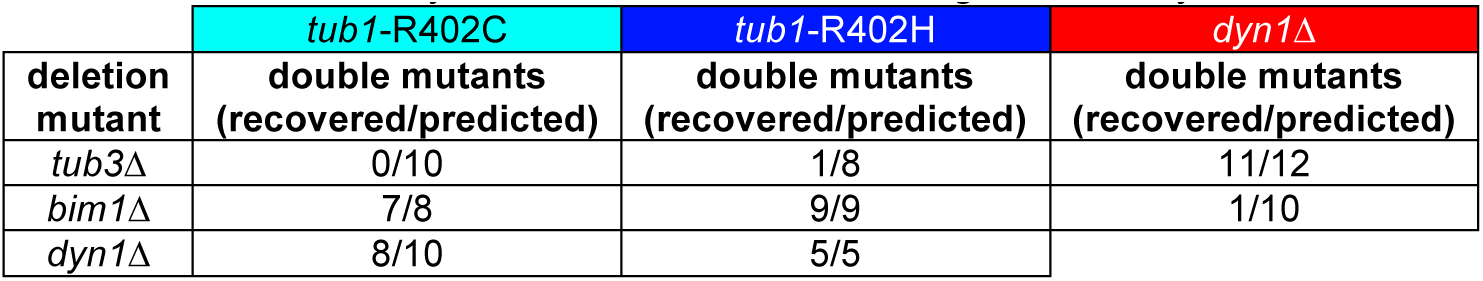
Survival of yeast double mutant tetrads generated by meiotic cross.

**Table S6. Yeast strains used in this study**

**Table S7:** Vectors used in neuronal assays

| Construct | Plasmid # | Promoter | Selection | Assay |
| --- | --- | --- | --- | --- |
| pCIG2 | pJM295 | CAG | Amp | In utero electroporation, ICC |
| pCIG2-Tuba1a(WT)-ires-GFP | pJM416 | CAG | Amp | In utero electroporation, ICC |
| pCIG2-Tuba1a(R402C)-ires-GFP | pJM418 | CAG | Amp | In utero electroporation, ICC |
| pCIG2-Tuba1a(R402H)-ires-GFP | pJM420 | CAG | Amp | In utero electroporation, ICC |
| pCIG2-Tuba1a(WT)-ires-GFP-MACF43 | pJM430 | CAG | Amp | Microtubule dynamics |
| pCIG2-Tuba1a(R402C)-ires-GFP-MACF43 | pJM432 | CAG | Amp | Microtubule dynamics |
| pCIG2-Tuba1a(R402H)-ires-GFP-MACF43 | pJM434 | CAG | Amp | Microtubule dynamics |
| pCIG2-Tuba1a(P263T)-ires-GFP-MACF43 | pJM514 | CAG | Amp | Microtubule dynamics |
| pCIG2-GFP-MACF43 | pJM436 | CAG | Amp | Microtubule dynamics |
| pCIG2-mNeonGreen-Lyso-20 | pJM452 | CAG | Amp | Axonal trafficking |
| pCIG2-Tuba1a(WT)-ires-mNeonGreen-Lyso-20 | pJM449 | CAG | Amp | Axonal trafficking |
| pCIG2-Tuba1a(RC)-ires-mNeonGreen-Lyso-20 | pJM450 | CAG | Amp | Axonal trafficking |
| pCIG2-Tuba1a(RH)-ires-mNeonGreen-Lyso-20 | pJM451 | CAG | Amp | Axonal trafficking |

**Table S8.**
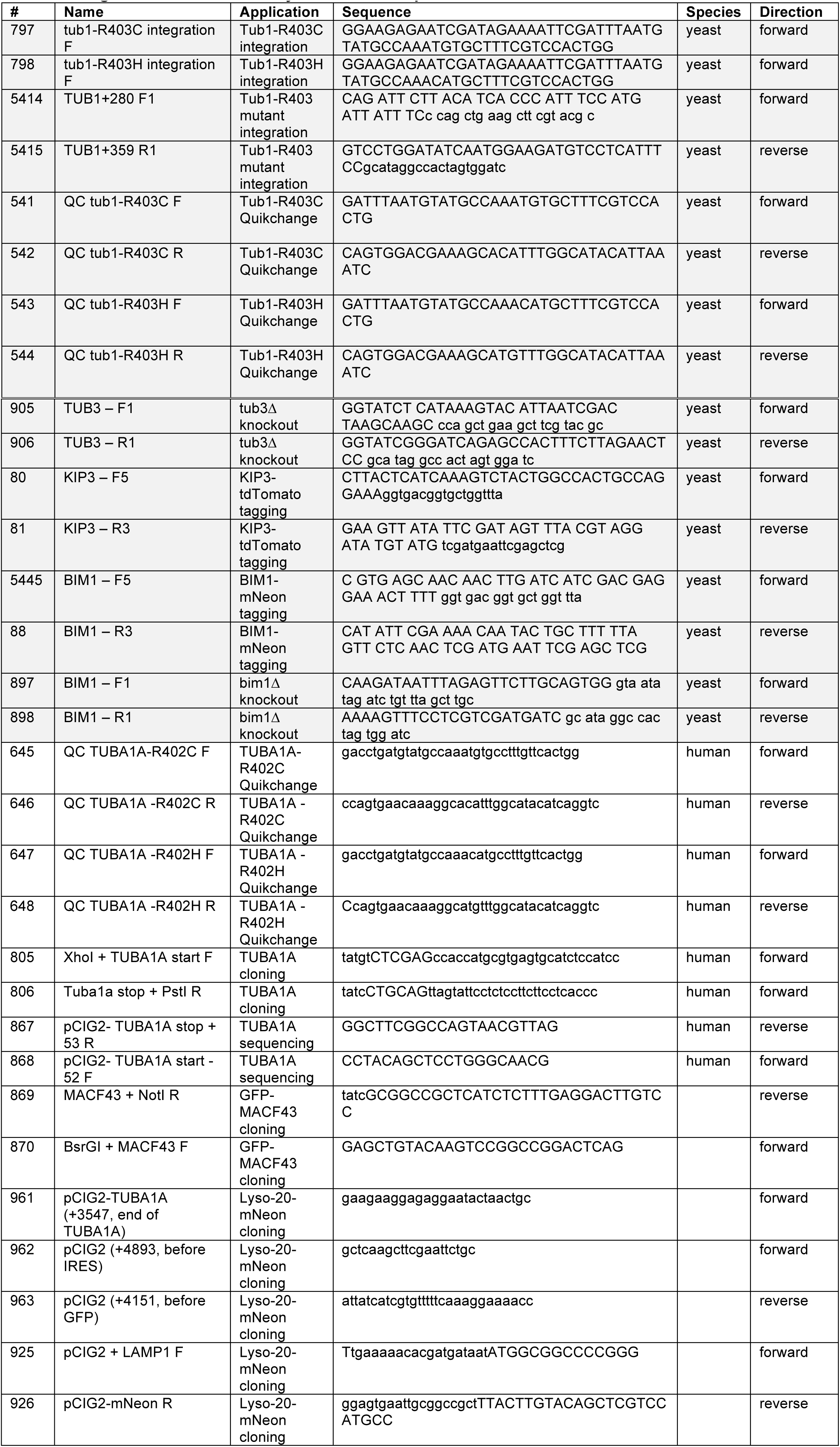
Oligos used in PCR-based yeast strain and plasmid construction

